# Pangenome structure and ecological adaptation in the *Klebsiella pneumoniae* species complex: insights from a geographically and time limited multi-habitat study

**DOI:** 10.64898/2025.12.11.693626

**Authors:** Jose F. Delgado-Blas, Elodie Barbier, Virginie Passet, Catherine Neuwirth, Sylvain Brisse, Carla Rodrigues, Pascal Piveteau

**Affiliations:** Institut Pasteur, Université Paris Cité, Biodiversity and Epidemiology of Bacterial Pathogens, Paris, France; AgroEcologie, UMR 1347, INRAE-UBE-IAD, Dijon, France; Laboratoire de Bactériologie, Centre Hospitalier Universitaire de Dijon, Dijon, France; INRAE, UR OPPALE, Rennes, France

## Abstract

*Klebsiella pneumoniae* species complex (KpSC) members inhabit distinct ecological habitats, yet the diversity and adaptation of environmental KpSC populations remain underexplored. We investigated KpSC transmission, pangenome structure and diversity, and functional gene enrichment across four distinct habitats in Burgundy, France over one year. In total, 664 environmental samples were collected from an organic vegetable farm (n=329), an organic cattle farm (n=304) and wastewater treatment plants (WWTP; n=31), alongside 47 clinical isolates. KpSC was detected in 22.4% of environmental samples, most commonly in WWTP (83.9%), followed by vegetable farm (27.1%) and cattle farm (11.2%). A total of 336 isolates were collected and whole-genome sequenced. *K. pneumoniae sensu stricto* (phylogroup Kp1) was predominant (76%), followed by *K. variicola* subsp. *variicola* (phylogroup Kp3) (21%). Genomic analyses revealed substantial novel diversity, especially in environmental habitats (67 novel STs), with limited cross transmission but clear local persistence. Over 90% of environmental isolates lacked acquired antimicrobial resistance genes, whereas half of the clinical isolates exhibited a multi-drug resistance profile. Comparative pangenome analyses showed Kp1 possessed a larger and more diverse pangenome than Kp3. Human-associated populations (clinical and WWTP) shared similar pangenome structures, although clinical isolates exhibited an expanded accessory genome. Cattle farm isolates had the most restricted and distinct pangenome while vegetable farm isolates displayed the largest total genomic repertoire. Functional enrichment analysis highlighted Kp3’s environmental adaptation via conserved functions linked to the phylogroup genetic background, such as metabolic and regulatory pathways, including nitrogen fixation genes. Contrarily, human-associated populations, especially Kp1 members, were enriched in acquired functions linked to the ecological context, including antimicrobial and metal resistance determinants and mobile genetic elements. These findings emphasize phylogroup- and ecological niche-driven pangenome contrasts in KpSC, contributing to explain the successful adaptation of Kp1 to multiple habitats, including human-related settings.

## Introduction

*Klebsiella pneumoniae* is considered one of the most threatening bacterial pathogens for public health worldwide, mainly due to the emergence and dissemination of multi-drug resistant clonal groups (CGs) (1,2). However, clinically relevant CGs are only a small fraction of the extensive *K. pneumoniae* diversity, which is largely composed of environmental populations (3). Besides, at the species level, *K. pneumoniae sensu stricto* (phylogroup Kp1) is one of the seven closely related phylogroups that form the *Klebsiella pneumoniae* species complex (KpSC) alongside *K. quasipneumoniae* subsp. *quasipneumoniae* (phylogroup Kp2) and subsp. *similipneumoniae* (phylogroup Kp4), *K. variicola* subsp. *variicola* (phylogroup Kp3) and subsp. *tropica* (phylogroup Kp5), *K. quasivariicola* (phylogroup Kp6), and *K. africana* (phylogroup Kp7) (4,5), but their diversity and ecology are poorly known.

KpSC members are versatile ubiquitous microorganisms (6). Their ecological habitats include the gastrointestinal tract of humans and animals, but also water, sediments, effluents, soil, plants, and some isolates were even characterised as plant growth promoters (7). The ecological diversity of KpSC is paralleled by its extensive metabolic versatility (8–10). However, adaptations of KpSC members to their various habitats, as well as the potential role of the environment as a reservoir for clinically relevant KpSC CGs and antimicrobial resistance (AMR) genes, remain poorly understood (11). Some studies have identified wastewater treatment plants (WWTPs) (12,13), farms (14), and food of animal origin and vegetables (15,16) as reservoirs facilitating the dissemination of clinically relevant KpSC strains and AMR genes, but transmission events between environmental and clinical habitats remain difficult to capture (3).

*K. pneumoniae* has a high capacity to collect, integrate and disseminate genetic material through multiple mobile genetic elements (MGEs), resulting in an open pangenome characterized by a large accessory genome (11,17). This genomic plasticity facilitates the rapid acquisition of genes involved in ecological adaptation, supporting the broad ecological range and persistence of *K. pneumoniae* as both an environmental organism and an opportunistic pathogen. AMR and virulence factors are key traits observed in clinical *K. pneumoniae* isolates, but they represent a small subset of the much larger adaptive genetic repertoire found in this bacterial species complex. Beyond that, the impact of ecological context on pangenome structure, as well as the influence of phylogroup-specific constraints on the dynamics of genetic gain and loss, are still unknown aspects (18). Consequently, our current understanding of KpSC ecological and genomic complexity is both limited and biased, as most available genomes come from clinical strains primarily selected based on AMR determinants, highlighting the need for further research within a One Health framework (3,19).

The present study aimed to characterize the KpSC population structure and transmission dynamics, as well as their pangenome structure and diversity across species and their ecological niche. We employed a geographically and temporally restricted sampling strategy, collecting samples over one year from two organic farms (vegetable and cattle), effluents and sludges from local WWTPs, and clinical isolates from the local hospital. Integrating extensive in-depth core- and pan-genome analyses with this comprehensive sampling strategy provides novel insights into how the phylogeny and ecological niche shape KpSC population diversity and environmental adaptation.

## Materials and Methods

### 1. Study design and sampling strategy

A geographically restricted, time-limited sampling strategy was carried out to generate a collection of KpSC isolates from multiple ecological origins. The Côte-d’Or department, in the Bourgogne-Franche-Comté region, including the geographical area around the French city of Dijon, was selected due to the availability of diverse habitats, including farms, WWTPs and hospital environments (**Figure 1A**). Samples were collected between July 2018 and July 2019. Throughout, we distinguish three distinct ecological concepts: i) ’habitat’, the broad spatial context of a bacterial population (*e.g.*, human, WWTP, human-associated, farm, environment-associated); ii) ’ecological source’, the specific matrix or substrate from which an isolate was recovered (*e.g.*, clinical infection, sludge, soil, water, root, leave, dung); and iii) ’ecological niche’, the combination of gene content, metabolic capabilities, and functional adaptations, that permit a bacterial population to thrive and persist within specific habitats (20,21).

**Figure 1.**
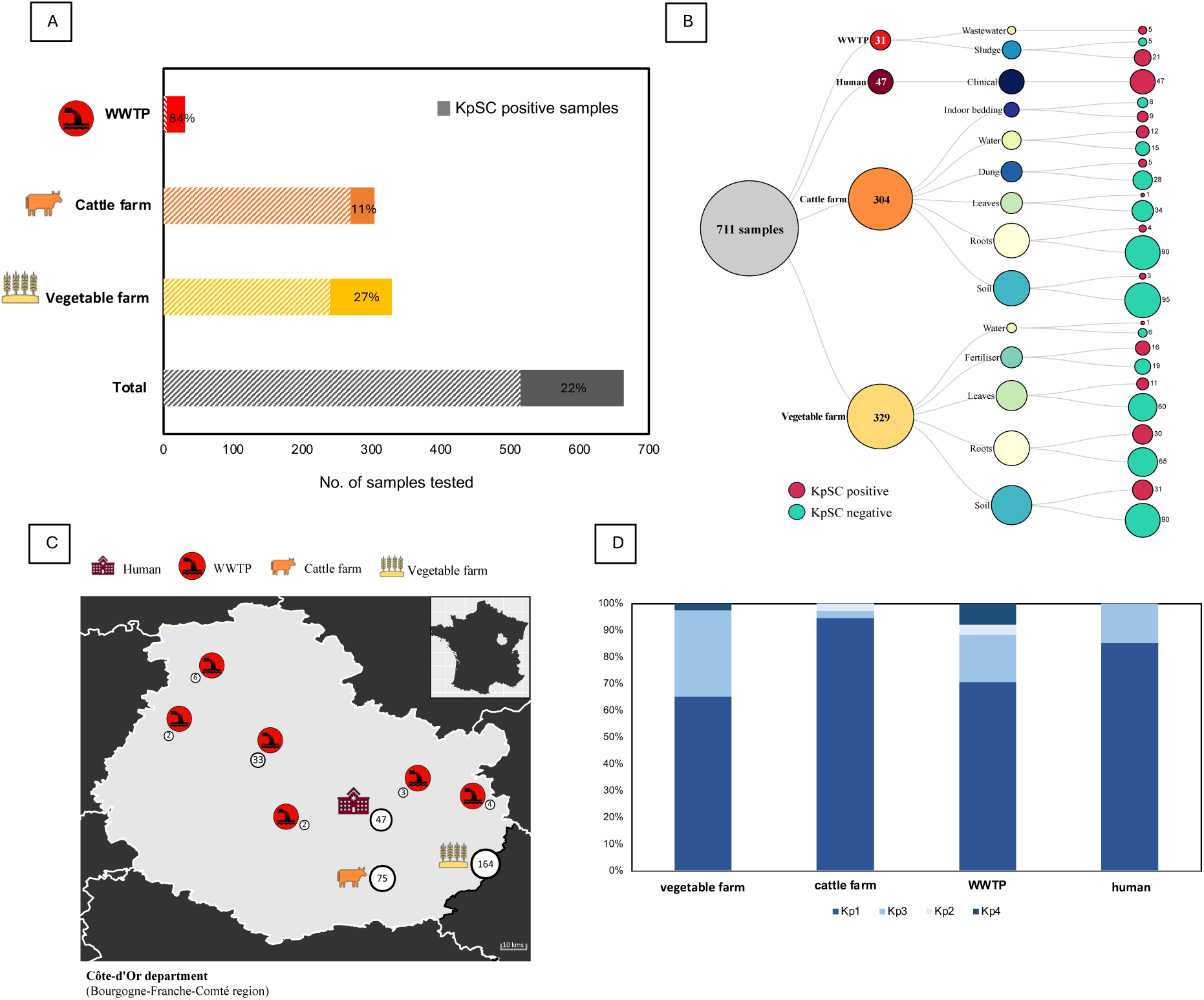
Distribution of samples and KpSC isolates from this study. **A:** Prevalence of KpSC positive samples globally and across the different habitats. **B:** Distribution of positive and negative samples according to habitat and ecological source (bottom legend). Circle sizes are proportional to the number of samples, which are indicated in/by the circles. **C:** Geographical and ecological distribution of KpSC isolates collected in the Côte-d’Or department. Habitat types are represented by symbols (top legend) and the number of KpSC isolates from each sampling location is indicated by circle numbers and sizes. **D:** Distribution of KpSC phylogroups across the different habitats.

One organic vegetable farm and one organic cattle farm were visited periodically to collect samples from the same plots. Vegetable farm samples (n=329) consisted in bulk soil (n=121), edible plants (parsley, basil, spinach, lettuce) and weeds. Root system (n=95) and leaves (n=71) were sampled from whole plants when present. Irrigation water (n=7) and organic fertilizer/compost (n=35) were also sampled. Cattle farm samples (n=304) consisted in bulk soil (n=98), pasture grass roots (n=94) and leaves (n=35), water (trough, well and river) (n=27), cow dung and winter indoor cow bedding (n=50). In total, 633 farm samples were collected and processed in the lab within 24h of sampling for detection and isolation of KpSC (**Figure 1B**). Besides, six local community WWTPs were sampled (n=31). Samples of 24h treated water (n=5) were provided by the “Laboratoire Départemental de la Côte d’Or”. Stored liquid activated sludge (n=24), dehydrated sludge (n=1) and composted sludge (n=1) samples were provided by the “Chambre d’agriculture de Côte d’Or” (**Figure 1B**). Finally, 47 unique clinical KpSC isolates collected during the same period were provided by the bacteriology laboratory of the main hospital in Dijon (CHU Dijon-Bourgogne). These clinical isolates were primarily from septicaemia patients, but also included urine, wound, respiratory and abscess samples.

### 2. Isolation and identification of KpSC

Isolation and identification of KpSC was performed according to Barbier *et al*. (22). Briefly, leaves and roots were cut and cleaned with sterile distilled water and transferred in 180 mL plastic containers. Ten-gram soil, dung, indoor bedding and sludge samples were weighed and also transferred into 180 mL plastic containers and then suspended in 90 mL of lysogeny broth (LB) supplemented with ampicillin (10 mg/L). Water samples (500ml) were filtered (0.25µm membrane) and the membrane filters were immersed in 20 mL of LB supplemented with ampicillin. All LB enrichments were incubated 24h at 30°C, vortexed, and 500 µL aliquots were centrifuged (5min, 5,800g) and washed with sterile water. The pellet was resuspended in 500 µL of sterile water and boiled for 10 min. Boiled enrichments were 10-fold diluted (1:10 and 1:100), and the dilutions were used as templates for ZKIR qPCR to detect KpSC positive enrichments (22).

KpSC-positive LB enrichments were streaked on Simmons citrate agar supplemented with 1% inositol (SCAI medium), and incubated for 48 hours at 37°C (23). Presumptive KpSC colonies (large, yellow, dome-shaped) were screened, with up to ten candidates purified and confirmed by MALDI-TOF MS. Up to three confirmed isolates per sample were selected for further analyses. Clinical non-redundant KpSC strains were isolated from Drigalski agar and identified by MALDI-TOF MS. All isolates were stored frozen at -70°C in glycerol stocks.

### 3. Whole-genome sequencing, genome assembly and typing analyses

Genomic DNA extraction of 337 selected KpSC isolates (**Table S1**) was automatically performed by MagNA Pure 96 (Roche Life Sciences, Sweden), following the manufacture’s guidelines. Genomic library was obtained using Nextera XT Prep Kit, and subsequent sequencing was carried out using the NextSeq 500 Illumina technology (Illumina, San Diego, USA), generating 2×150 paired-end reads. Illumina reads were processed and *de-novo* assembled with *fq2dna* tool (https://gitlab.pasteur.fr/GIPhy/fq2dna). Resulting assemblies were quality controlled (QC) using *contig_info* tool (https://gitlab.pasteur.fr/GIPhy/contig_info) and assembly metrics were verified to comply with the KlebNET-GSP consortium’s QC criteria (https://bigsdb.pasteur.fr/klebsiella/genomes-quality-criteria/).

Genotype classifications based on multi-locus sequence typing (MLST) (24), core-genome MLST (cgMLST) (25) and Life Identification Number (LIN) codes (5) were determined using the *Klebsiella* BIGSdb-Pasteur web application (BIGSdb-Kp) (https://bigsdb.pasteur.fr/klebsiella/). Genetic diversity of KpSC across the four habitats was assessed using rarefaction analysis and Simpson’s Index of Diversity (SID) based on unique ST and cgST counts. Rarefaction curves were generated with the *vegan* R package to compare richness standardized by sample size. SID and 95% confidence intervals were calculated via bootstrapping, and pairwise habitat differences were tested using permutation with multiple testing correction. BIGSdb-Kp and Kleborate v2 (26) were used to determine AMR, virulence, and heavy metal tolerance genes content, and to predict the capsular (K) and O-antigen (LPS) serotypes. Likewise, plasmid replicons were identified using PlasmidFinder (27).

### 4. Pangenome structure and diversity

Pangenome analyses were performed with Roary v3.12 (28), using as input genomes annotated with Prokka v1.12 (29). Gene clusters were defined as core genes when present in ≥90% of the genomes and using a blastP identity cutoff of 80%. Single-nucleotide polymorphisms (SNPs) were extracted from the resulting core-genome alignment by SNP-sites (30) and used to generate a maximum likelihood phylogenetic tree with IQ-TREE v1.6.11 applying a GTR+F model (31). Tree visualization and edition were carried out with iTOL v6.8.1 (32).

Simultaneously, another specialized pangenome approach developed for bacterial ecological studies was performed to integrate, analyse, and compare ecological niche- and phylogroup-specific KpSC genomic contents. The anvi’o platform v8 (33) was applied following the workflow for microbial pangenomics. Briefly, an anvi’o database was created with the KpSC genomes, including gene-calling data from Prodigal (34) and functional characterization from NCBI’s Cluster of Orthologous Groups (COG) database (35). Pangenome analyses using optimized parameters (see supplementary material, **Figure S12**) were performed subsequently either for the total KpSC population, or for Kp1 and Kp3 independently, or for the habitat-specific subpopulations (*i.e.*, clinical, WWTP, cattle farm and market garden farm). We applied a *minbit* score of 0.5 to filter meaningful amino acid sequence matches (two sequences share a bit score that is at least 50% of the bit score obtained when either sequence is aligned to itself); a Markov clustering (MCL) inflation of 8 or 10 (for total and phylogroup-specific populations, respectively), which defines the sensitivity of amino acid cluster identification and controls the granularity of clusters (higher values produce more, smaller clusters than lower values, leading to finer distinctions between groups); and a 95% gene presence threshold to define the core genome. Pangenome profiles were visualized and edited using the anvi’o interactive interface. Accumulation curves of total pangenomes, core-genomes and accessory genomes, were assessed, using the *panplots* v1.0 R package (https://doi.org/10.5281/zenodo.6408803) with R v4.3.2, and visualized with *ggplot2* v3.4.4 R package (36) in RStudio v2023.12.0+369 (37). Rarefied pangenome and core-genome sizes of the different populations were estimated by gene cluster accumulation of the less represented group over 100 iterations and statistically evaluated by T-test. Gene presence/absence data, including singleton clusters, were analysed using Jaccard and Manhattan distance metrics, as previously described (38,39). Mobilome-associated gene clusters (e.g., prophages, transposons, plasmids) were similarly analysed.

Core-genome clusters (*i.e.*, single-copy gene-clusters present in at least 95% of the genomes) were extracted, concatenated and used to generate an approximately-maximum-likelihood phylogenetic tree (*anvi-get-sequences-for-gene-clusters* and *anvi-gen-phylogenomic-tree* programs, respectively). The resulting tree was used to determine the distance to the most recent common ancestor (MRCA) (see supplementary material). Phylogenetic trees, including extracted phylogroup/niche subtrees, were also used to calculate pairwise patristic distances using the *cophenetic* function in the *ape* R package. Additionally, DNA sequences from the core genome (single-copy gene clusters with 1.0 geometric homogeneity or 0-gap alignments) were extracted to assess nucleotide diversity (π) using the *nuc.div* function in the *pegas* R package (40).

Genome metrics (including genome length, number of gene clusters, and %GC content) were extracted and compared across the total KpSC population, Kp1 and Kp3, and their ecological niche-specific subpopulations. Results were visualized with *ggplot2* and statistically analysed (see supplementary material). Additionally, pangenome structure, pangenome diversity, core-genome distances, and genome metrics were compared to assess their interactions within the entire KpSC population and the Kp1 and Kp3 subpopulations, with statistical significance evaluated (see supplementary material).

### 5. Phylogenomic and enrichment analyses

Functional enrichment analyses were performed using anvi’o based on COG functional categories (41) (see supplementary material). Functions significantly associated with a specific KpSC phylogroup and ecological niche (adjusted *q*-value<0.05) were labelled as ‘phylogenetically enriched functions’ while those linked to particular ecological niches, but not to KpSC phylogroups, were considered ‘ecologically enriched functions’. Results were visualized and edited with the anvi’o interactive interface. The diversity of enriched functions across phylogroups and ecological niches was assessed using Jaccard distances from curated presence/absence profiles, followed by visualization and statistical analysis, as done for pangenome and mobilome diversity.

## Results

### 1. KpSC members distribution in different habitats and ecological sources

Over the 12 months sampling period, KpSC strains were detected in 149 out of the 664 environmental samples (22.4%) (**Figure 1A**). The overall ecological distribution of KpSC positive environmental samples was 83.9% from WWTP (26 samples), 27.1% from the organic vegetable farm (89 samples), and 11.2% from the organic cattle farm (34 samples) (**Figure 1A**). In terms of source type in vegetable farm, KpSC presence was the highest in organic fertilizers (16 samples, 45.7%), followed by roots (30 samples, 31.6%) and soil (31 samples, 25.6%); whereas in the cattle farm, occurrence of KpSC was the highest in indoor bedding (9 samples, 52.9%) and water (12 samples, 44.4%) (**Figure 1B**).

In total, 289 KpSC environmental isolates were collected; 239 KpSC isolates were recovered from the farms (164 from the vegetable farm and 75 from the cattle farm), and 50 isolates were obtained from WWTP samples (**Figure 1C, Table S1**). Kp1 accounted for 73.7% of the environmental isolates (214/289), being the predominant phylogroup in all habitats and sources sampled (**Figure 1D**). The remaining environmental isolates were identified as Kp3 (64 isolates, 22%), Kp4 (7 isolates, 2.4%) and Kp2 (4 isolates, 1.4%). While most Kp3 isolates (60.6%) were recovered from soil, roots, and leaves sampled across both farms, underscoring the soil dwelling nature of this phylogroup (**Figure 1D**), a statistically significant association between the vegetable farm and the presence of Kp3 was also depicted (*p* < 0.00001). The vast majority of the clinical isolates were Kp1 (n=40, 85.1%), distantly followed by Kp3 (n=7, 14.9%) (**Figures 1B**

### 2. Population structure and diversity of KpSC across different habitats

Based on the 7-loci MLST scheme, the KpSC population was distributed into 156 different sequence-types (STs), including 67 (42.9%) newly defined STs, numbered non consecutively between ST4041 and ST4972 (**Figure 2**, Microreact link: https://microreact.org/project/total-kpsc-population, BIGSdb project id 54 at https://bigsdb.pasteur.fr/cgi–bin/bigsdb/bigsdb.pl?db=pubmlst_klebsiella_isolates). Most of these novel STs, among both Kp1 and Kp3 isolates, originated from environmental sources (86.5%), especially from the vegetable farm (53.7%). Among all STs, ST200 was the most common overall (n=21), identified in clinical, vegetable farm, and cattle farm samples. The recognized SL307 high-risk clone member ST307 was the most prevalent (n=7) among clinical isolates, and it was also detected in one WWTP sample. In cattle farm samples, ST4439 was the most frequent one (n=11), while ST4422 predominated in the vegetable farm (n=11).

**Figure 2.**
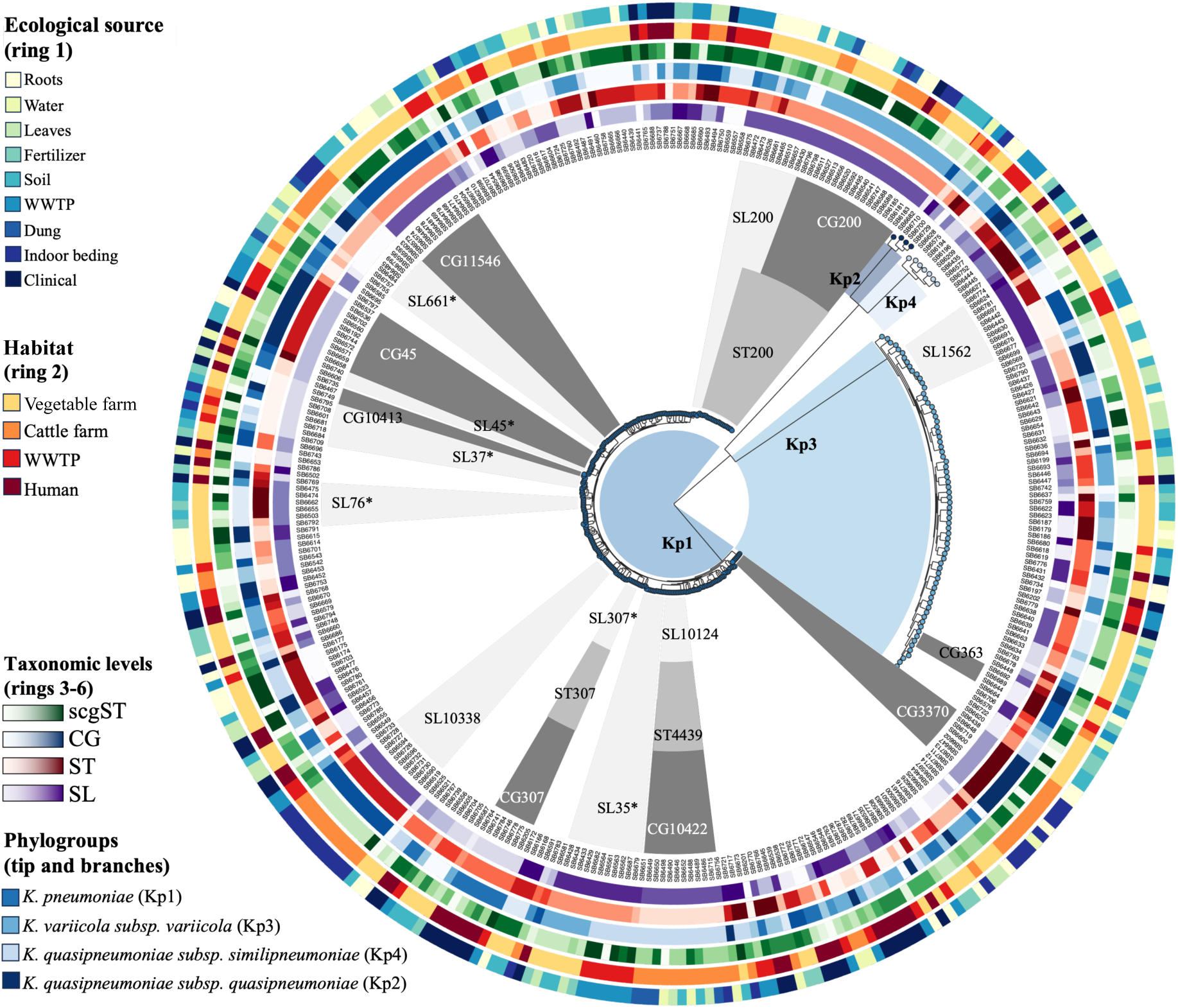
Phylogenetic and ecological structure of KpSC population. Tips of the phylogenetic tree are coloured according to the KpSC phylogroup, as well as the branch shaded sectors (bottom-left legend). Ring 1 (most outer ring) and ring 2 are coloured regarding the ecological source and habitat, respectively (top-left legends). Rings 3 to 6 (most inner ring) are alternately coloured according to decreasing discriminatory taxonomic levels of the KpSC cgMLST scheme as follows: core-genome sequence type (scgST, ring 3), clonal group (CG, ring 4), sequence type (ST, ring 5) and sublineage (SL, ring 6). Grey-shaded sectors between the tree tips and the rings show the most representative SLs, STs and CGs, including those considered high-risk SLs (*). Interactive visualization at Microreact website: https://microreact.org/project/total-kpsc-population

Based on the 629-loci cgMLST scheme, 248 cgSTs were identified, of which 211 (85.1%) were novel, highlighting again the diversity of KpSC population (**Figure 2**). These cgSTs were distributed in 143 sublineages (SLs) and, in concordance with MLST results, SL200 was the most prevalent SL in the total population (26 isolates; 7.7%). Indeed, SL200 was detected in clinical samples as well as in multiple sources of the cattle and vegetable farms, followed by SL45 (11 isolates; 3.3%) also present in multiple habitats (human, vegetable farm and WWTP) and SL37 (3.0%, all habitats). Notably, 19% of the SLs were classified as high-risk SLs (42), especially the human-associated SLs from clinical and WWTP samples (40%), but also SLs from the vegetable and cattle farms (12% from each environment).

We assessed the genetic diversity of KpSC across the four habitats using rarefaction analysis and SID at both ST and cgST resolutions (**Figure S1**). Rarefaction curves demonstrated that clinical, WWTP and vegetable farm habitats exhibited the highest expected richness across standardized sample sizes, with largely overlapping curves, while the cattle farm showed markedly lower richness. Consistently, diversity estimates based on SID showed that vegetable farm isolates had the highest diversity at the ST level (SID = 0.970, 95% CI: 0.962–0.976), followed by WWTP (SID = 0.944, 0.922–0.956) and clinical (SID = 0.939, 0.911–0.955) sources. Cattle farm isolates had the lowest diversity (SID = 0.909, 0.878–0.930). Given the higher cgST resolution, diversity was higher than for STs, but ranking was similar: vegetable farm was the highest (SID = 0.982, 0.979–0.985) and cattle farm the lowest (SID = 0.945, 0.924–0.960). Pairwise permutation tests confirmed that ST diversity in cattle farm was statistically lower than in clinical (*p* = 0.012), WWTP (*p* = 0.0021), and vegetable farm (*p* < 0.001). Similarly, cgST diversity was significantly lower in cattle farm relative to vegetable farm (*p* < 0.001), while other comparisons were not statistically significant after multiple testing correction.

### 3. Transmission dynamics of KpSC within and between habitats

To detect potential recent transmission events both across and within habitats, and to assess persistence over time within habitats, we defined common genotypes using a two-step approach. First, genotypes were considered common if they showed ≤ 4 mismatches out of 629 loci in the KpSC cgMLST scheme (5,42). This was complemented by a second criterion: a cutoff of ≤21 SNPs in the core-genome alignment obtained through Roary (43) (**Table S2**).

Only two cases of common genotypes based on the first criterion (cgMLST mismatches) were depicted between habitats. The first involved two CG10413-ST37 isolates (from spinach leaves and a clinical blood culture, in 2019) which differed by 3 cgMLST alleles only, but 58 core-genome SNPs. The second involved two CG290-ST290 isolates (from indoor cattle bedding and soil from a garden pea field), differing by a single cgMLST allele but 95 SNPs. Neither case met the second criterion (≤21 SNPs), not supporting recent cross-habitat transmission or the presence of a shared source for these genotypes.

Within habitats, we observed strong evidence of persistence and recent transmission, characterized by minimal genomic differences using the double criteria, particularly in the farms (**Table S2**). At the cattle farm and vegetable farm, almost half (37% and 41%, respectively) of the total isolates belonged to common genotypes (between 0 and 4 cgMLST allele differences and 0-21 SNP distances), indicating colonization with limited microevolution (*i.e.*, recent transmission) (**Table S2**). Genotypes were shared either across different ecological sources over time, within the same source over time, or between different components of the same source (*e.g.*, a plant or pasture) at one or multiple time points (**Table S2**). These findings demonstrate the sustained presence and localized transmission of KpSC genotypes within individual farms or habitats. In contrast, common genotypes were less frequently observed in the clinical setting and WWTPs. In the clinical setting, three cases of human infection involved the same genotype: CG10234-ST70 (Kp1), CG10802-ST589 (Kp1), and the well-known CG15-ST15 (Kp1). All these isolates differed by 5 to 11 SNPs, likely reflecting nosocomial transmission (**Table S2**). In only one instance, two samples from different WWTPs collected on the same day shared the same genotype (CG10741-ST4045, Kp3), with a difference of only 3 SNPs.

### 4. Surface antigens, virulome, resistome and plasmidome profiles

Capsular locus characterization identified 137 *wzi* alleles and 70 different KL types. The KL type was not predictable in 28.8% of the isolates, especially from the vegetable farm (58 isolates, 35.4%). Similarly, O-antigens were also more diverse in the vegetable farm, with up to 11 different types. Clinical and WWTP populations presented similar O-antigen profiles and O1 was the most frequent type. However, O2 was the most frequent one in fertilizer samples, O3a in leaves and O3b in roots and indoor bedding (**Figure 3, Figure S2**).

**Figure 3.**
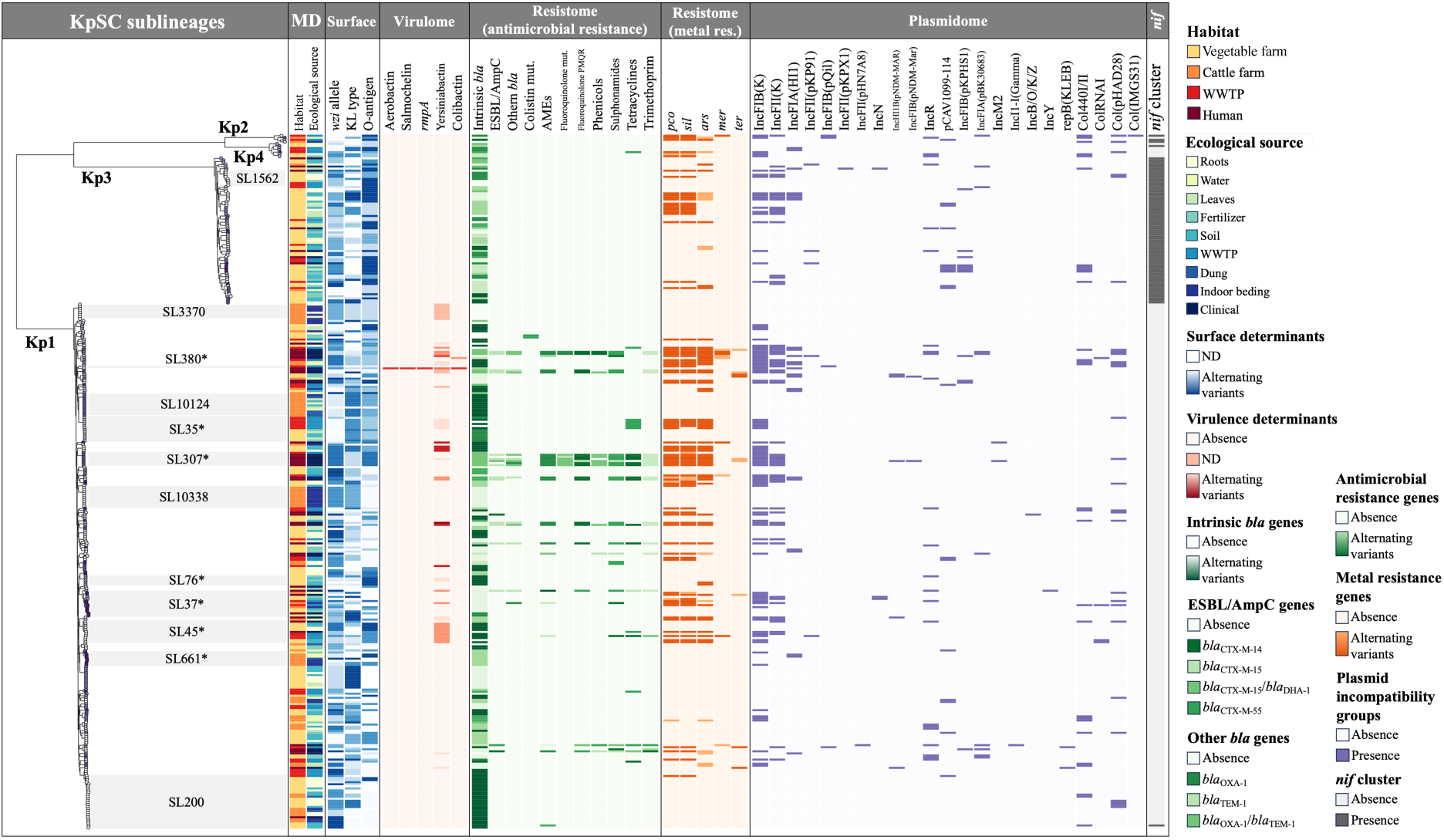
Genetic profiles of total KpSC population. Tips of the phylogenetic tree are alternately coloured according to sublineage (SL), and grey-shaded sectors show the most representative SLs, including those considered high-risk SLs (*). Metadata (MD) columns are coloured according to the habitat (column 1) and ecological source (column 2) (see top-right legend). The presence and variety of genetic determinants belonging to surface antigens (columns 3 to 5), virulence factors (columns 6 to 10), antimicrobial resistance genes (columns 11 to 21), metal resistance genes (columns 22 to 26), plasmid incompatibility groups (columns 23 to 49) and *nif* genes (column 50) are displayed as indicated in the different right-side legends. Complete profiles with genes and variants for all genetic determinants are included in Table S1. Genes and variants of ESBL/AmpC (column 12) and other β-lactamases (column 13) are specified due to their clinical relevance. ND: Non determined. AMEs: aminoglycoside-modifying enzymes. Colistin mut.: chromosomal mutations conferring resistance to colistin. Fluoroquinolone mut.: chromosomal mutations conferring resistance to fluoroquinolones. Fluoroquinolones PMQR: plasmid-mediated quinolone resistance.

The yersiniabactin gene cluster was the only virulence gene cluster found across all habitats, and it was frequent in clinical isolates (38.3%) with seven distinct lineages identified (**Figure 3**). Lineage *ybt*10, linked to ICEKp4 and high-risk SL45, was widespread, while *ybt*9 carried on ICEKp3 was detected in a range of high-risk SLs which varied according to the habitat, notably SL35 in WWTPs. One hypervirulent SL380 clinical isolate carried the genes encoding the five virulence factors aerobactin, salmochelin, capsule, yersiniabactin, and colibactin (**Figure 3, Table S1**).

The distribution of antimicrobial resistance genes (ARGs) across sources is shown **Figure 3**. Most environmental isolates (94.5%) did not carry extra ARGs beyond the intrinsic beta-lactamase genes *bla*_SHV,_ *bla*_LEN_ or *bla*_OKP,_ according to their phylogroup. In sharp contrast, half of the clinical population exhibited multidrug-resistant profiles with several resistance genes, notably the ESBL gene *bla*_CTX-M-15_ (36.1%), *qnrB1* (29.7%), *sul2* and *drf14* (36.1% both), conferring resistance to extended-spectrum cephalosporins, fluoroquinolones, sulfonamides and trimethoprim, respectively (**Figure 3**, **Figure S3**). In WWTP isolates, acquired ARGs were detected in 19.6% of samples, primarily *tet* genes associated with tetracycline resistance.

Regarding the heavy metal tolerance genes, *pco* (copper), *sil* (silver) and *ars* (arsenic) were strongly associated with clinical (60.0%) and WWTP (47.1%) sources (**Figure 3**, **Figure S3**). The presence of ARGs was directly linked to the plasmid content, which depended on the habitat. Thus, no plasmid replicons were identified in two thirds of the vegetable farm (65.2%) and the cattle farm (62.7%) populations, whereas in human and WWTP habitats the percentage of isolates with at least one plasmid replicon was notably high (87.2% and 64.7%, respectively) (**Figure 3, Table S1**). IncFIB(K) was the most frequent replicon, found in 94 isolates of different phylogroups and ecological sources, but mostly in clinical (68.1%) and WWTP (39.2%) populations. It usually cohabited with other replicons such as IncFII(K). Despite similar plasmidome and metal resistance profiles found in human-associated populations, clinical and WWTP isolates showed different antimicrobial resistance patterns.

### 5. Pangenome structure of KpSC population

Based on anvi’o, the KpSC pangenome was composed of 17,688 gene clusters, including singletons, of which only 3,017 (17.1%) were defined as part of the core genome (**Figure S4**). These results were highly concordant with those from Roary (3,946/22,005 gene clusters; 17.9% core-genome). At the phylogroup level, the Kp1 pangenome was made of 14,708 out of the total 17,688 gene clusters, divided in a core-genome of 3,289 genes (22.4%) and a large accessory genome (11,419 gene clusters). Kp3 had a more reduced pangenome (10,773 gene clusters), with a larger core genome (4,015, 37.3%) and less accessory genes (6,758 genes). Due to the limited number of Kp2 and Kp4 isolates, the pangenome structure of these phylogroups could not be confidently evaluated, since pangenome analyses are largely dependent on population representativeness. Therefore, Kp2 and Kp4 genomes were not included in subsequent analyses.

To account for differences in sample size between phylogroups, accumulation curves and rarefaction analyses were performed. The results (**Figure 4**) showed that Kp1 and Kp3 exhibited nearly identical accumulation patterns for total gene clusters (**Figure 4A**), with rarefied pangenome sizes of 10,875 and 10,773 gene clusters, respectively. Despite this similarity, the Kp1 pangenome was statistically larger than that of Kp3 (t = 2.79, *p* = 0.0067). While the rarefied core-genome of Kp1 was significantly smaller than that of Kp3 (t = –47.10, *p* < 10⁻⁵⁵) (**Figure 4B**), a significantly greater number of unique accessory genes was found in Kp1 than in Kp3 (t = 14.54, *p* = 0) (**Figure 4C**). The pangenome curves thus reflects a more conserved global genetic repertoire in Kp3, while Kp1 appears to be composed of a more diverse population with specific genetic contents.

**Figure 4.**
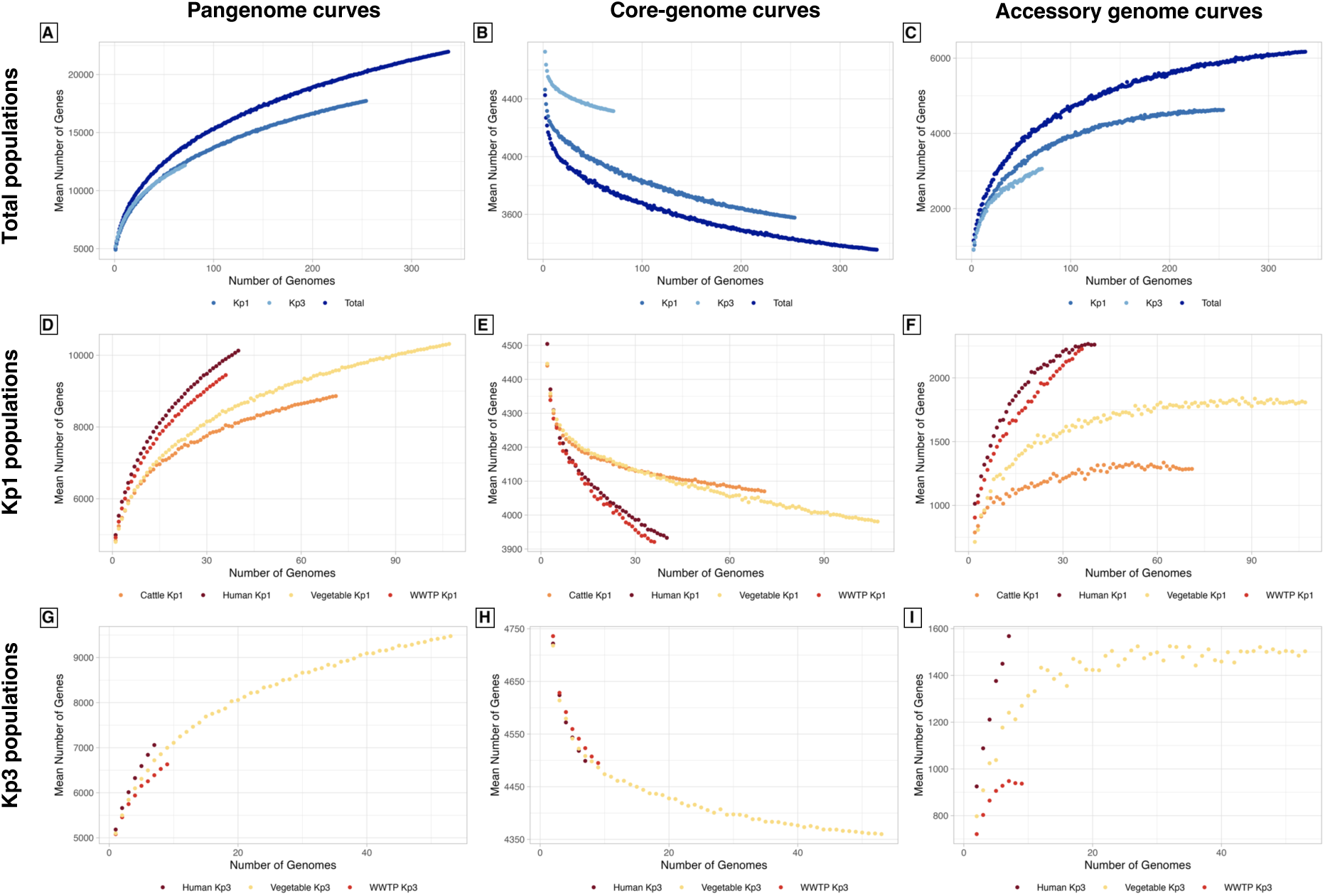
Pangenome, core-genome and accessory genome accumulation curves of KpSC populations. Each panel shows the accumulated gene clusters (*Y* axis) according to the increasing number of genomes (*X* axis). Legend for each panel is at the right side. **A:** Pangenome accumulation curves of total KpSC, Kp1 and Kp3 populations. **B:** Core-genome accumulation curves of total KpSC, Kp1 and Kp3 populations. **C:** Accessory genome accumulation curves of total KpSC, Kp1 and Kp3 populations. **D:** Pangenome accumulation curves of Kp1 populations according to habitats. **E:** Core-genome accumulation curves of Kp1 populations according to habitats. **F:** Accessory genome accumulation curves of Kp1 populations according to habitats. **G:** Pangenome accumulation curves of Kp3 populations according to habitats. **H:** Core-genome accumulation curves of Kp3 populations according to habitats. **I:** Accessory genome accumulation curves of Kp3 populations according to habitats. Note data for cattle farm Kp3 population are not presented due to the low number of Kp3 genomes from this reservoir (n=2).

When comparing the pangenome size across habitats, KpSC populations from clinical source (3,606 core-genes/11,397 gene clusters) and WWTP (3,569 core-genes/11,023 gene clusters) showed highly similar sizes (**Figures S5A and S5B**). The cattle farm population possessed a more limited pangenome (3,792 core-genes/9,894 gene clusters) (**Figure S5C**), whereas the vegetable farm population exhibited a broader pangenome (3,463 core-genes/13,133 gene clusters) (**Figure S5D**).

To account for a possible phylogroup effect, we next compared pangenomes across habitats for each phylogroup. Similar core-genome-pangenome correlations were observed in Kp1 and Kp3 populations (z = 0.733, *p* = 0.232; **Figure 5A**). As expected, these correlations were significantly negative for both phylogroups, since the pangenome increase led to a core-genome reduction. Clinical populations of Kp1 and Kp3 exhibited a larger rarefied pangenome caused by a faster increase of pangenome clusters than the other populations (**Figure 4D, 4G**), linked again to a rapid reduction of core-genome clusters (**Figures 4E and 4H**) and the increase of accessory gene clusters (**Figures 4F and 4I**). The Kp1 and Kp3 clinical rarefied pangenomes were not statistically different between them (t=-2.48, *p*=0.05), but the clinical Kp1 rarefied core-genome was significantly lower than the Kp3 one (t=-16.45, *p*<10^-5^), showing the same pangenome trends than the total Kp1 and Kp3 populations (**Figure 5A).** Clinical pangenome curves were closely followed by the WWTP population in the case of Kp1 and the vegetable farm population in the case of Kp3 (**Figures 4D-4I**), but rarefied pangenome clusters were significantly smaller compared to the clinical populations (t=-2.69, *p*=0.019, and t=-3.42, *p*=0.014, respectively) (**Figure 5A)**.

**Figure 5.**
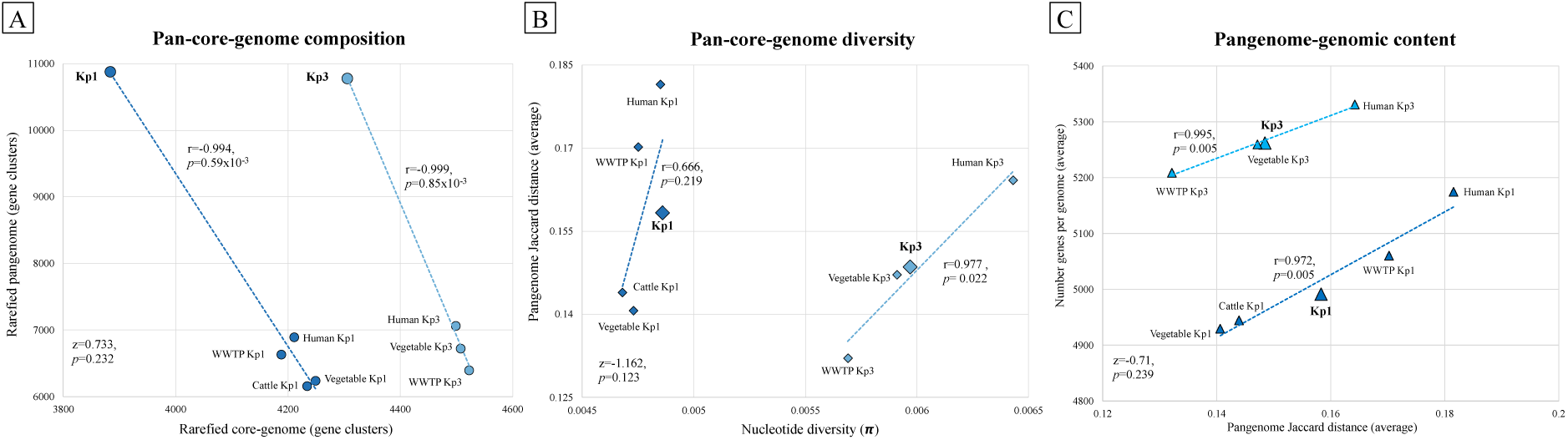
Pan-core-whole-genome interactions of total and habitat-specific Kp1 and Kp3 populations. Linear fits for Kp1 and Kp3 populations are shown by blue dashed lines, indicating the Pearson’s r and *p* values for each population correlation. Fisher’s z values for differences of correlations, and their corresponding *p* values, are indicated in the bottom-left corner of each panel. **A:** Association between the rarefied pangenome and rarefied core-genome. **B:** Association between the nucleotide diversity (π) and pangenome Jaccard distance. **C:** Association between the pangenome Jaccard distance and number of genes per genome.

In terms of genomic features (**Figure S6**), the genome length and genetic content were highly correlated, as expected, in both Kp1 (r=0.999, *p*=0.42×10^-4^) and Kp3 (r=0.997, *p*=0.26×10^-2^) populations, but Kp3 genomes were larger (∼300 kb) and thus harboured a higher genetic content (∼250 more genes) than Kp1 genomes (**Figure S6A**). Regarding the GC content, it exhibited a strong negative correlation with the gene number in both phylogroups (Kp1: r=-0.993, *p*=0.63×10^-3^; Kp3: r=0.990, *p*=0.98×10^-2^). Generally, GC content of Kp1 populations was lower than that of Kp3 populations (**Figure S6B)**, consistent with AT-richness of the overall accessory genome (44). At intra-phylogroup level, clinical populations of both phylogroups presented longer genomes with more genes and lower GC contents than the other populations, followed again by WWTP for Kp1 and organic vegetable farm for Kp3 populations (**Figures S6A and S6B**).

### 6. Pangenome diversity across KpSC populations

Pangenome diversity analyses based on Jaccard distances revealed that total Kp1 population exhibited more diverse gene cluster profiles than Kp3 population (t=14.55, p<2.2×10^-16^) (**Figure S7A**). Clinical populations from both phylogroups possessed the most diversified genetic contents, followed by WWTP Kp1 population and vegetable farm Kp3 population. However, the pangenome diversity of different ecological Kp3 populations was more homogeneous compared to Kp1 populations (**Figure S7A)**.

To determine if the faster evolution of Kp1 pangenome is due to its larger accessory genome size or if the pangenome diversity simply reflects a more genetically diverse population, we compared pangenome Jaccard distances of total and habitat-specific Kp1 and Kp3 populations with their respective core-genome nucleotide diversities (**Figure 5B**). Surprisingly, Kp1 populations exhibited lower nucleotide diversity than Kp3 and, therefore, the Kp1 broader pangenome diversity was not the result of a more diverse population. While total and habitat-specific Kp3 populations showed a strong positive correlation between pangenome diversity and nucleotide diversity (r = 0.977, *p* = 0.022), pangenome diversity in Kp1 was not significantly correlated with nucleotide diversity (r = 0.666, *p* = 0.219) (**Figure 5B**). There was a significant correlation for both phylogroups between the number of genes per genome and pangenome diversity (Kp1: r=0.972, *p*=0.005; Kp3: r=0.995, *p*=0.005) (**Figure 5C**). Nevertheless, Kp1 populations presented larger pangenome diversities than Kp3 ones, although their genomes carried a lower genetic content.

At intra-phylogroup level, clinical populations presented higher pangenome and nucleotide diversities (**Figure 5B**). Likewise, to test if a larger nucleotide diversity was related with a longer evolutionary divergence, the nucleotide diversities were compared with the MRCA distances (**Figure S6C**). No significant correlation was found neither for Kp1 (r=-0.076, *p*=0.903) nor Kp3 (r=-0.60, *p*=0.394) populations, indicating that the higher nucleotide diversity of Kp3 was not the result of a deeper phylogenetic structure than Kp1 population. However, clinical populations possessed the highest nucleotide diversity in the shortest phylogenetic distance to the MRCA, pointing out the population diversification effect of this ecological source (**Figure S6C**).

To address whether Kp1 pangenome diversity is the result of a larger phylogenetic divergence between its members, we examined the relationship between pangenome diversity and phylogenetic distance. As expected, the overall KpSC pangenome diversity showed a strong positive correlation with patristic distance or, in other words, genetic content divergence between two genomes increased with phylogenetic distance (r = 0.68, *p* < 0.001) (**Figure S8A**). Interestingly, this relationship differed between phylogroups: for the total Kp1 population, there was a statistically significant positive correlation (r = 0.11, *p* < 0.001). In contrast, the Kp3 population exhibited a small but statistically significant negative correlation (r = –0.06, *p* = 0.003), suggesting that pangenome diversity within Kp3 was not increasing according to phylogenetic divergence (**Figure S8B**). These findings indicate that the larger accessory genome observed in Kp1 may be more driven by population segregation and lineage specialization than by accelerated genome evolution.

Regarding the mobilome diversities, considering the presence/absence profiles of gene clusters related with prophage, transposon and plasmid elements, Kp3 population carried a more diverse genetic content than Kp1 population (t=-5.73, *p*=1.11×10^-8^) **(Figure S7B**). However, when looking at the ecological niche-specific populations, there were no statistical differences, except for the higher mobilome diversity of the Kp3 population found in the vegetable farm compared to the Kp1 population from the same habitat and the WWTP Kp1 population compared to the other Kp1 populations (**Figure S7B**).

### 7. Phylogenetically and ecologically enriched functions in KpSC populations

With the aim of identifying genetic clusters potentially linked to phylogroup- and/or ecological niche-specific populations, subsequent functional enrichment analyses were performed based on pangenome profiles. Both phylogenetically and ecologically enriched functions, including COG categories and adjusted *q*-values for phylogroup and ecological niche assignation, are included in Tables S3A and S3B, respectively. Firstly, 97 functions were significantly enriched in specific KpSC phylogroups from specific ecological niches (‘phylogenetically enriched functions’) (**Figure 6, Figure S9**). Most of these functions were associated with metabolism (58.8%) and regulatory processes (19.6%), and revealed the strongest phylogroup-ecological niche connection between Kp3 members and vegetable farm (**Figure 6A, S9A**). Remarkably, multiple genes of the nitrogen fixation *nif* operon were identified as key traits of Kp3 populations from the vegetable farm (**Figure 3, Table S3A**). Contrarily, no phylogenetically conserved functions were found in the Kp1 population isolated from the organic vegetable farm, since the functions they carried, including virulence and structural traits, were indiscriminately related with all other habitats (**Figure 6A, Figure S9A**).

**Figure 6.**
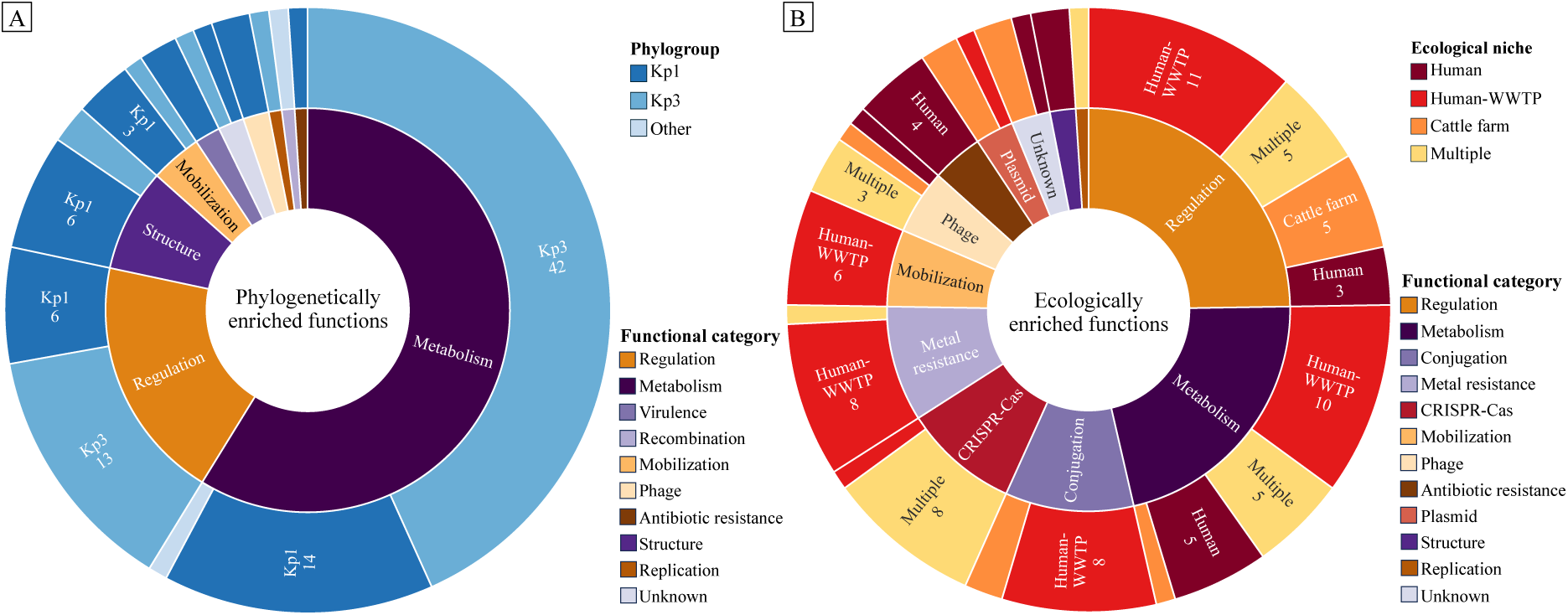
Schematic visualization of phylogenetically and ecologically enriched functions. **A:** Phylogenetically enriched functions. Inner ring indicates the functional category of all phylogenetically enriched functions (right legend). Outer ring shows the phylogroup in which the functional category is enriched with the number of enriched functions related to that category. **B:** Ecologically enriched functions. Inner ring indicates the functional category of all ecologically enriched functions (right legend). Outer ring shows the ecological niches in which the functional category is enriched with the number of enriched functions related to that category.

On the other hand, 97 distinct functions were exclusively related with the ecological niche regardless of the phylogroup (‘ecologically enriched functions’) (**Table S3B)**. Most of them were assigned to regulation (24.7%) and metabolic processes (21.6%), and, to a lesser extent, other functions such as conjugation (10.3%), metal resistance (9.3%) and mobilization (6.2%) (**Figure 6B, Figure S9B**). Some functions were significantly related with clinical isolates, especially aminoglycoside and β-lactam resistance mechanisms. Most ‘ecologically enriched functions’ were shared by clinical and WWTP populations, consistent with their similar pangenome profiles. Among them, we found genes encoding resistance to the heavy metals tellurium (*ter*), copper (*cop*) and arsenic (*ars*), as well as conjugation structures (type IV secretion system), and mobile genetic elements (Tn*3*, IS*26* or IS5 among others) (**Figure 6B, Figure S9B**). The cattle farm population displayed specific functions such as conjugation-related elements and toxin-antitoxin systems, highlighting special biotic interactions in this ecological context. None of the ‘ecologically enriched functions’ was exclusively associated to the vegetable farm.

We then compared phylogenetically and ecologically enriched functional profiles to assess their diversity across Kp1 and Kp3 populations. Regarding phylogenetically enriched functions, functional profiles of Kp1 and Kp3 populations were drastically distinct, as expected, due to their deep evolutionary divergence (**Figure S10A**). In contrast, Kp1 and Kp3 members within the same ecological niche shared ecologically enriched functional profiles (**Figure S10B**), indicating convergent or recent ecological adaptation despite phylogenetic differences.

When focusing on functional diversity of these enriched functions, the Kp1 population exhibited significantly greater overall diversity of phylogroup-enriched functions compared to Kp3 (t = 8.98, *p* = 2.2×10⁻¹⁶), as well as across different ecological niche-specific subpopulations (**Figure S11A**). Moreover, these phylogroup-enriched profiles were significantly more diverse among all Kp1 niche-specific populations, while niche-specific Kp3 populations showed less diverse, more homogeneous phylogroup-enriched functional profiles. Similarly, the diversity of ecologically enriched functions was significantly higher in Kp1 overall (t = 113.33, *p* = 2.2×10⁻¹⁶). However, unlike phylogroup-enriched functions, ecological niche-specific Kp3 populations exhibited equal or even greater diversity of ecologically enriched functions than Kp1 (**Figure S11B**). Additionally, while Kp1 populations showed significant differences in ecologically enriched functions across habitats, Kp3 niche-specific populations did not differ significantly, maintaining similar phylogenetic and ecological functional profiles regardless of their origin. This contrast indicates that functional diversity of Kp1 members is more dependent on the ecological conditions, whereas Kp3 members exhibit a conserved functional profile regardless of their environmental context.

## Discussion

The genetic diversity and adaptability of KpSC populations, especially *K. pneumoniae sensu stricto* (Kp1), have been extensively demonstrated in clinical contexts, where this species is recognized as part of the ESKAPEE group, largely driven by its genetic ability in acquiring antimicrobial resistance (AMR) genes (45). Similarly, in environmental contexts, especially soil and aquatic systems, KpSC members are widely spread and recurrently identified (3,6,46–48). However, knowledge about the diversity, transmission and adaptability of KpSC populations in natural and anthropogenic environments remains limited. Current challenges constraint our understanding of environmental KpSC ecology, such as the lack of investigation of multiple habitats beyond clinical settings, the underrepresentation of antimicrobial susceptible populations in KpSC strain collections, and the scarcity of accurate environmental genomic data and metadata. In complement to previous attempts (3,19,49), our study addresses these gaps by systematically sampling and characterizing a spatiotemporally defined KpSC population spanning human-, animal-, and environment-associated habitats, with unbiased selection criteria with regards to AMR phenotype. Moreover, this work further explores the extent of KpSC diversity and adaptability in terms of genomic content and ecological context. KpSC prevalence varied considerably across environmental habitats, with the highest occurrence in WWTP samples (83.9%), followed by vegetable farm (27.1%) and cattle farm (11.2%). Phylogroup analysis revealed Kp1 as the predominant group across all habitats (representing 73.7% of environmental isolates), while Kp3 was significantly associated with environmental sources such as soil and roots, particularly from samples collected in the vegetable farm. These results support the ubiquity of KpSC, especially Kp1 populations, and confirm previous findings (3,49).

The detection of a substantial proportion of novel STs and cgSTs within the studied KpSC population highlighted a remarkable diversity among the sampled habitats, especially the organic vegetable farm. This extensive hidden diversity reflects an unexplored and complex genetic landscape beyond clinical settings. Diversity metrics demonstrated that vegetable farm, WWTP, and clinical populations harboured higher genetic diversity than the cattle farm population, suggesting that the cattle farm represents a more bounded system, likely reflecting the colonization and transmission of a locally conserved, bovine-specific population. This ecological heterogeneity between habitats aligns with the concept of ‘ecological compartmentalization’ previously observed in other studies (3,19,49), whereby distinct habitats foster ecological adaptation even within CGs. Indeed, fine-scale genetic analyses using cgMLST and SNP distances applied to STs found in different habitats (*e.g.*, ST200 from clinical and environmental sources), revealed limited evidence of cross-habitat transmission events. In contrast, within-habitat comparisons, particularly within farm environments, demonstrated persistent colonization and local transmission of closely related KpSC genotypes over time. Indeed, the recurrent detection of genotypes with minimal genetic variation from cattle (*e.g.*, CG10422-ST4439) and vegetable (*e.g.*, CG11546-ST4422) farms evidenced sustained habitat-specific residency. This pattern was less pronounced in clinical and WWTP settings, where fewer recent transmission clusters were detected. These findings suggest that persistence of environmental strains and intra-habitat transmission may be more frequent than expected, whereas human-related habitats favour frequent population replacement caused by transient strong selective pressures (38).

Analysis of AMR profiles also revealed highly contrasted patterns across habitats. Notably, over 90% of WWTPs and farms isolates lacked acquired AMR genes, while half of the clinical population exhibited a multi-drug resistance profile, indicating that environmental strains are highly distinct from clinical populations. Besides, these findings suggest, on the one hand, a limited spillover from clinical to environmental habitats and, on the other hand, a minor role of environmental populations as direct reservoirs of AMR strains, as also previously observed in other studies (3). Besides, heavy metal resistance genes and plasmid replicons were mostly absent in the farm habitats but largely enriched in human-associated populations (WWTP and clinical), even if these populations differed in their AMR genetic contents. Thus, the ‘ecological compartmentalisation’ effect not only influenced the KpSC population structure, but also their plasmidic and AMR genetic contents, as previously described in several works (3,49,50). However, while all these studies observed infrequent cross-habitat bacterial and genetic flux, there is currently little mechanistic insight into what genomic factors may influence the ecological adaptation of these distinct KpSC populations to specific environments. Moreover, analysis of large genomic collections spanning broad geographical regions and timeframes or the focus on specific phylogroups (mainly Kp1) may hamper significant eco-genetic understanding due to unexpected confounding factors such as specific geographic and temporal characteristics or limited phylogenetic representativeness, respectively.

Given its spatiotemporal restriction, the assembled collection of genomes was highly suitable to address broad ecological and evolutionary questions within KpSC: Do the pangenome structure and diversity depend on the phylogroup and/or the ecological niche? What are the genomic mechanisms behind distinct pangenomes and how do they influence ecological adaptation? Are there specific genetic clusters that provide ecologically adaptive functions and/or fitness-relevant features? Do these functions/features and their diversity depend on the phylogroup or are they linked to the ecological source? And ultimately, what is the role of these pangenome and functional dynamics in the broad-range adaptability of Kp1?

Comparison of Kp1 and Kp3 pangenome structures underscored distinct evolutionary dynamics between these phylogroups. Although both phylogroups shared similar total pangenome sizes, Kp1 displayed a larger accessory genome and smaller core genome, indicating higher genomic plasticity and potential specialization among its lineages. Conversely, Kp3 exhibited a larger, more conserved core genome concomitant with a smaller accessory genome, suggesting long-term stable adaptation to their habitat (51). Besides, genomic features further differentiated phylogroup-specific populations, since Kp3 genomes were larger and exhibited a higher GC content than those of Kp1 members. On the other hand, when comparing the pangenome structure across ecological niche-specific populations, Kp1 and Kp3 clinical isolates showed accelerated expansion in accessory genes and concomitant core genome reduction, consistent with ongoing adaptive diversification driven by clinical selective pressures. Additionally, clinical and WWTP populations shared similar pangenome profiles, confirming the common genomic tendencies of human-related KpSC populations. The cattle farm supported a reduced and more stable pangenome, in line with the population diversity results, while vegetable farm populations harboured the broadest pangenomes, likely reflecting the ecological heterogeneity and genetic exchanges associated with the high strain diversity and persistence within this habitat (52). The comparison of genomic features between ecological niche-specific populations revealed that clinical isolates across both phylogroups tend towards longer genomes with lower GC content, a hallmark of recent accessory gene acquisition (44).

Surprisingly, Kp1 populations exhibited broader pangenome diversity despite lower nucleotide diversity across the core-genome compared to Kp3, suggesting that accessory genome expansion in Kp1 is more strongly influenced by ecological specialization. This was supported by positive correlations between pangenome diversity and patristic distances (phylogenetic divergence) in Kp1, contrasting with a weak negative correlation in Kp3, highlighting different evolutionary trajectories dependent on the interaction between phylogroup and environmental conditions. Altogether, this could be the result of a long divergent evolution, where Kp1 members adapted to human and animal related environments, leading to the reduction and conservation of the individual genome (53), but increasing the population genomic pool; whereas Kp3 members experienced a genome increase and diversification process (54), leading to a population pangenome conservation adapted to their habitat. Nevertheless, at intra-phylogroup level, both Kp1 and Kp3 clinical isolates presented a higher nucleotide diversity compared to the other populations, as well as larger and more diverse genomes and pangenomes, demonstrating the impact of ecological adaptation to clinical settings on pangenome expansion.

Concordantly with pangenome dynamics, enriched functions differed according to phylogroup and ecological niche. Phylogenetically enriched functions were strongly associated with Kp3 strains isolated from the vegetable farm. They included predominantly functions related to metabolic and regulatory pathways, including the nitrogen fixation *(nif*) operon, previously linked to plant endophytic relationships (11,51,55). In contrast, distinct phylogenetic functions could not be evidenced in Kp1 isolates from the organic vegetable farm, suggesting a less specialized environmental association. Ecologically enriched functions, regardless of phylogroup, included MGEs, heavy-metal resistance determinants, and AMR genes. These functions were notably enriched in clinical and WWTP populations, consistent with human-associated selective pressures (52). The cattle farm population exhibited unique enrichments in toxin-antitoxin systems and conjugation-related genes, pointing again at intra-habitat transmission and adaptation. Moreover, the diversity of these functional traits can be shaped by long-term evolutionary processes reflected in conserved phylogenetic-specific enrichments, or by more recent transient adaptations linked to ecological niches. In general, pangenome and enriched functional diversities followed similar inter-niche dynamics, where Kp1 divergence was explained by the reduction of the core-genomes and, therefore, the loss of phylogenetically enriched functions; whereas the Kp3 populations maintained more uniform pangenomes, which resulted in a higher content of phylogenetically related functions (51). Besides, when comparing the functional diversity across phylogroups, total Kp1 population demonstrated significantly higher diversity of both phylogenetically and ecologically enriched functions. This reflects a dynamic evolutionary landscape shaped by selective pressures and a large accessory genomic pool, key for its broad-range adaptability and its success in colonizing human-related contexts. In contrast, Kp3 populations showed conserved functional profiles regardless of their ecological niche, which is indicative of stable adaptation over extended evolutionary timescales.

This work presented several limitations. First, the geographically and time-limited sampling performed in several habitats is advantageous for ecologically driven pangenome analyses, but it should be extended across species and habitats in larger spatiotemporal contexts. Likewise, the underrepresentation (notably Kp2 and Kp4) or absence (Kp5-7) of certain phylogroups in the strain collection reduced comprehensive KpSC resolution, although this reflects their natural rarity. Additionally, nearly one-third of gene clusters lacked functional annotation, underscoring the lack of genetic investigation of environmental KpSC strains and the paucity of experimental validation. Likewise, the mechanistic role and ecological relevance of adaptive functions will need phenotypic confirmation through laboratory assays. However, the identification of previously known associations, such as the *nif* operon in environmental Kp3 populations, the AMR genes in clinical populations or the metal resistance genes in human-related populations, supports the methodology used and reinforces the adaptive potential of other identified phylogenetically and ecologically enriched functions. Despite these limitations, the integration of ecological and genomic data presented here, coupled with the comprehensive sampling strategy and in-depth pangenome analyses, revealed clear compartmentalization of KpSC populations shaped by their distinct pangenome structure and diversity. The conserved functional repertoire of Kp3 reflects its long-term adaptation to environmental contexts, whereas the broad accessory genome and functional diversity of Kp1 underpins its ecological expansion into multiple habitats, including human-associated settings. Therefore, the core genome reduction and the resulting functional diversification of a species compared with other non-pathogenic, closely related species could be indicative of increased adaptive potential and clinical risk.

## Data availability

Genome assemblies, quality control data, metadata and typing results are available at the *Klebsiella*-BIGSdb Pasteur website under the project id 54: https://bigsdb.pasteur.fr/cgi-bin/bigsdb/bigsdb.pl?db=pubmlst_klebsiella_isolates&page=project&project_id=54. Accession codes for all KpSC isolates and complete genetic profiles of surface antigens, virulence factors, antimicrobial resistance genes, metal resistance genes, plasmid incompatibility groups and *nif* genes are shown in Table S1. Graphical visualizations of provenance and phylogenetic data of the total KpSC population are available at Microreact website under the link: https://microreact.org/project/total-kpsc-population.

## Authors Contributions

Conceptualization: JFDB, CR, EB, PP and SB. Coordination: CR and SB. Environmental sampling strategy and wet-lab analysis and validation: EB and PP. Clinical sampling: EB and CN. Genomic sequencing and data curation: VP and CR. Genotyping analyses: JFDB and CR. Pangenome analyses, statistical inference and data visualisation: JFDB. Writing original draft: JFDB and CR. All authors contributed to writing and editing the manuscript and reviewed the final version. Funding Acquisition: SB.

## Acknowledgments

The authors acknowledge the Institut Pasteur teams for the curation and maintenance of BIGSdb-Pasteur databases at http://bigsdb.pasteur.fr/. We also thank the Mutualized Platform for Microbiology (P2M) for the genomic sequencing of the isolates using Illumina technology. This work used the computational and storage services provided by the IT Department at Institut Pasteur.

