## Supplementary Methods and Figures for "Pangenome structure and ecological adaptation in the *Klebsiella pneumoniae* species complex: insights from a geographically and time limited multi-habitat study"

##### Optimization of anvi'o parameters for pangenome analyses

For total KpSC, Kp1 and Kp3 population, habitat-specific subpopulations pangenome analyses, different combinations of *minbit* scores (0.3-0.8) and MCL inflations (8-10) were evaluated, selecting the aforementioned values as the optimal for gene cluster stabilization, capturing the diversity of non-related clusters and avoiding the over-segregation of closely related ones (**Figure S12**).

##### Distance to the most recent common ancestor (MRCA)

The core-genome phylogenetic tree generated using anvi'o was used to determine the distance to the most recent common ancestor (MRCA) of the total KpSC population, the Kp1 and Kp3 populations and the habitat-specific Kp1 and Kp3 populations. Briefly, the node of the MRCA for each population was identified with the *findMRCA* function of the *phytools* v2.1-1 R package(34), then the subtree for each population was extracted with the *extract.clade* function of the *ape* v5.7-1 R package(35), and the average distance to the MRCA of each of the population subtrees was calculated with the *nodeHeights* functions of *phytools* R package.

##### Statistical analyses for pangenome analyses

Statistical significance of Jaccard and Manhattan distances between Kp1 and Kp3 total and habitat-specific populations were assessed using T-test, whereas the statistical significance of all population-population distances was calculated by ANOVA and Tukey's HSD tests, considering the equality of variances and the normality assumption of the different groups, applying the Levene's test and the Shapiro-Wilk test, respectively. These statistical analyses were carried out with the *stats* v4.3.2 package integrated in R and graphically visualized with violin plots generated with *ggplot2* R package.

Statistical analyses of genome metrics' (*i.e.*, genome length, number of gene clusters and %GC content) differences between populations were performed as previously described for pangenome diversity results and, when any of the groups did not meet the assumption of normality criteria, the statistical significance was evaluated by performing Kruskal-Wallis test followed by *post-hoc* Mann Whitney U test, applying the Bonferroni correction. These analyses were performed with the *stats* and *MultNonParam* v1.3.9(37) R packages.

Pangenome diversity, pan-core-genome composition, core-genome distances, and genome metric values were compared to evaluate the pan-core-whole-genome interactions within the entire KpSC population as well as the distinct Kp1 and Kp3 subpopulations and the statistical significance was assessed. Firstly, patristic and pangenome Jaccard distance matrixes were pairwise combined, graphically visualized with *ggplot2* R package, and statistically analyzed applying the Spearman's rank correlation coefficient. Subsequently, average values of different distances and metrics were compared as follows: rarefied pangenome *vs.* rarefied core-genome, MRCA distance *vs.* nucleotide diversity, nucleotide diversity *vs.* pangenome Jaccard distance, pangenome Jaccard distance *vs.* number of genes per genome, and number of genes per genome *vs.* GC content. Then, these comparisons were statistically evaluated using the Pearson's correlation coefficient. Correlation coefficients were calculated with the *cor* function of the stats R package. Lastly, results from Spearman's and Pearson's coefficients from the different Kp1 and Kp3 populations were statistically compared applying the Fisher's *z*-test for differences of correlations in two independent samples with the *diffcor.two* function of the *diffcor* v0.8.2 R package(38).

**Functional enrichment analyses using anvi'o**

Each gene cluster from the pangenome was associated with a COG functional annotation, and the enrichment analyses based on the Rao test were independently performed for the total population considering the different KpSC species and ecological subpopulations. The resulting differential functions were filtered to retrieve significant hits (adjusted $q$-value<0.05; FDR-adjusted $p$-value for the Rao test), compared to pGWAS results, manually curated and categorized in general functional classes.

### Supplementary Figures

#### “Pangenome structure and ecological adaptation in the *Klebsiella pneumoniae* species complex: insights from a geographically and time limited multi-habitat study”

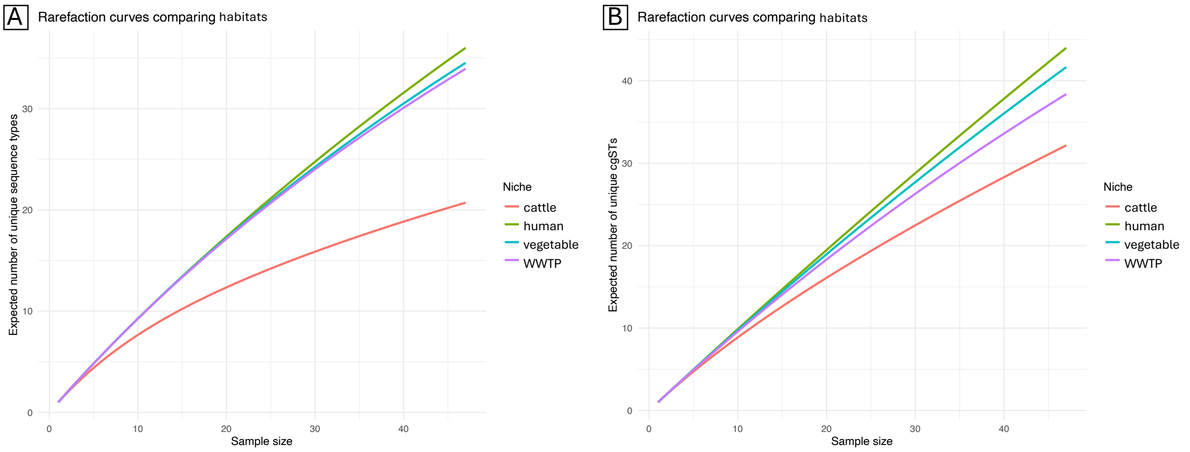


**Figure S1.** Population diversity of KpSC across the four habitats. Rarefaction curves were generated for each ecological niche to compare cluster richness (*Y*-axis), standardized by sample size (*X*-axis). The right panel legends indicate niche categories. **A:** Genetic diversity based on sequence types (STs). **B:** Genetic diversity based on core genome sequence types (cgSTs).

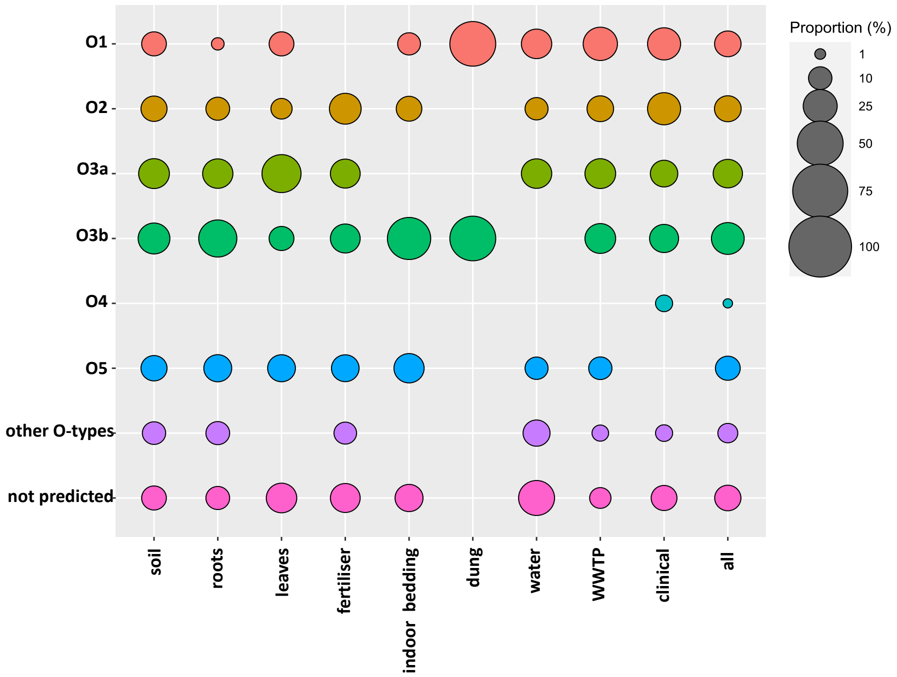


**Figure S2.** Distribution of O-antigen variants across the KpSC population. Different O-antigen variants are displayed in the y-axis and indicated by different colors. Ecological sources (sample types) and the total KpSC population are indicated in the x-axis. Size of then circles represent the proportion (%) of the O-antigen variant in each ecological source according to the top-right legend.

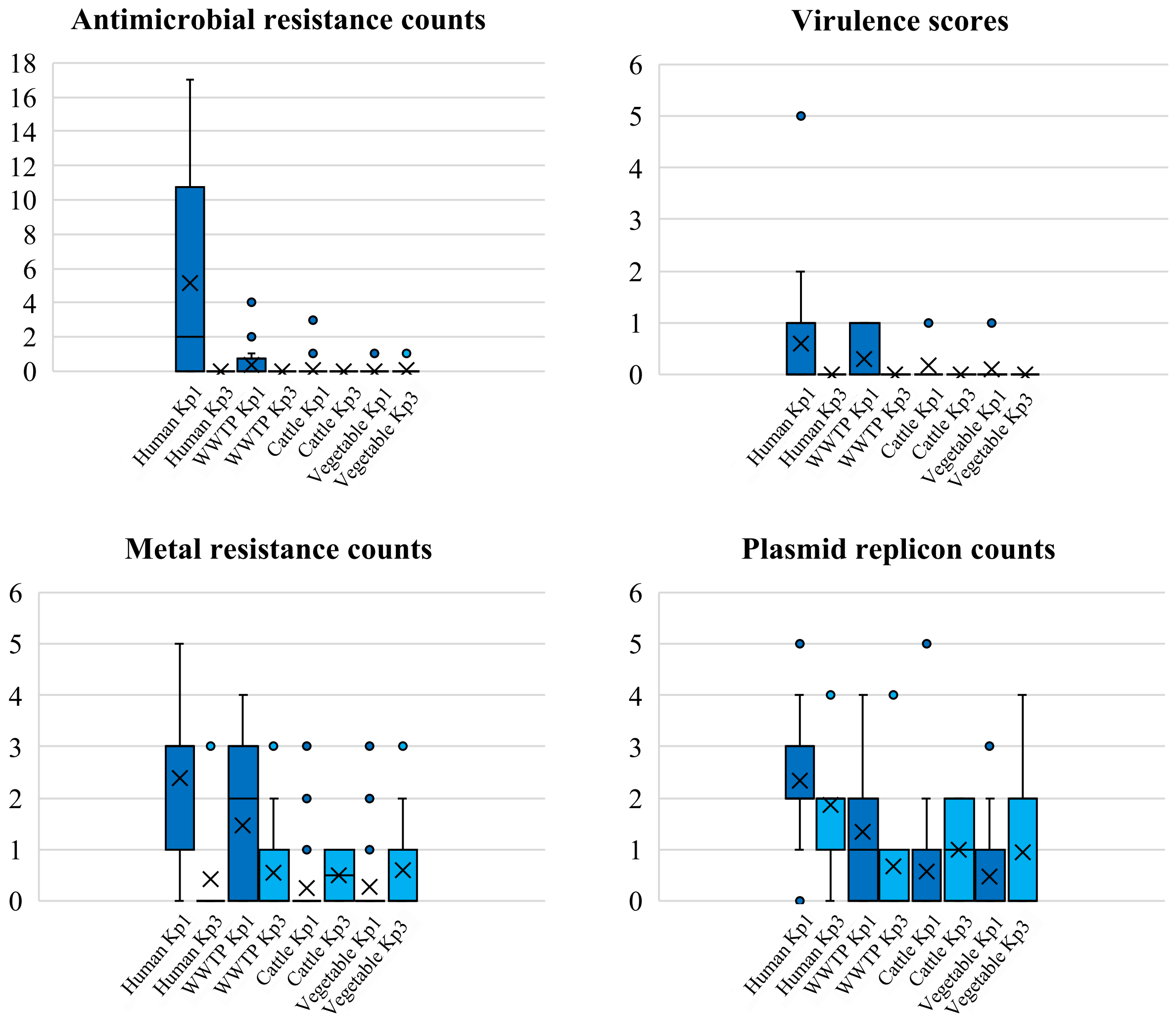


**Figure S3.** Distribution of genetic traits in the KpSC population according to phylogroup and habitat. Panels show boxplots by phylogroup (Kp1, dark blue; Kp3, light blue) and habitat (*X*-axis labels) for: i) antimicrobial resistance genes (Kleborate), ii) virulence scores (Kleborate), iii) metal resistance genes (Kleborate), and iv) plasmid replicons (PlasmidFinder). Boxes denote the median (line), the mean (X) and interquartile range (IQR), with points representing outliers.

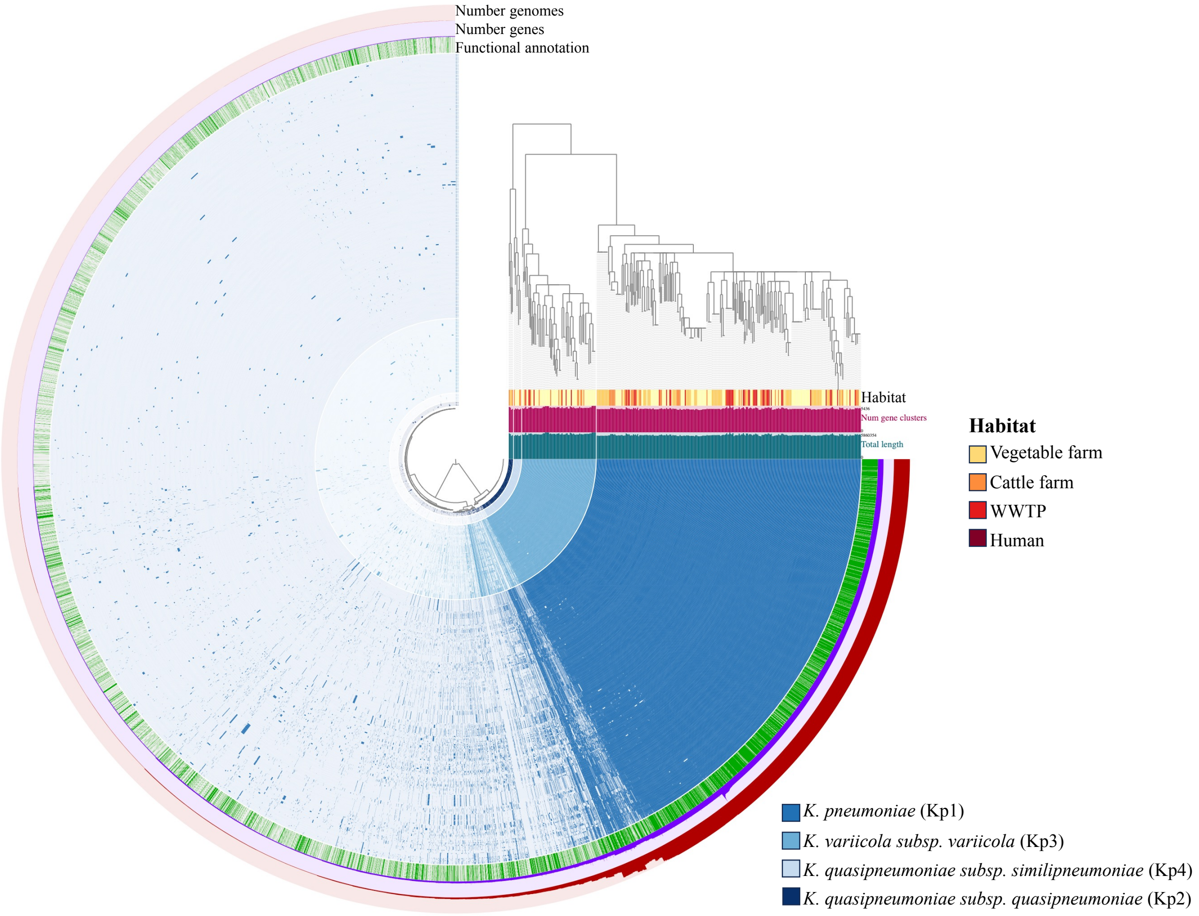


**Figure S4.** Pangenome structure of total KpSC population. Blue concentric rings display the gene cluster presence/absence profiles of all KpSC genomes. Blue color of the ring represents the KpSC phylogroup (bottom-right legend), whereas the blue tone indicates the presence (darker color) and absence (lighter color) of all gene clusters in the genome. The genomes are ordered according to the core-genome SNP tree of the total KpSC population (top-right corner). Between the tree and the rings, from bottom to top, the total genome length (dark green bar plots), the number of genes per genome (maroon bar plots), and the habitat (central right legend) are indicated for each genome. The tree in the center of the concentric rings represents the presence/absence distribution of each gene cluster across all genomes. The three most outer rings show, from inside to outside, the assignation of gene clusters to a COG category (dark green for assigned COG, light green for unknown COG), the number of genes in each gene cluster (purple ring), and the number of genomes possessing each gene cluster (red ring).

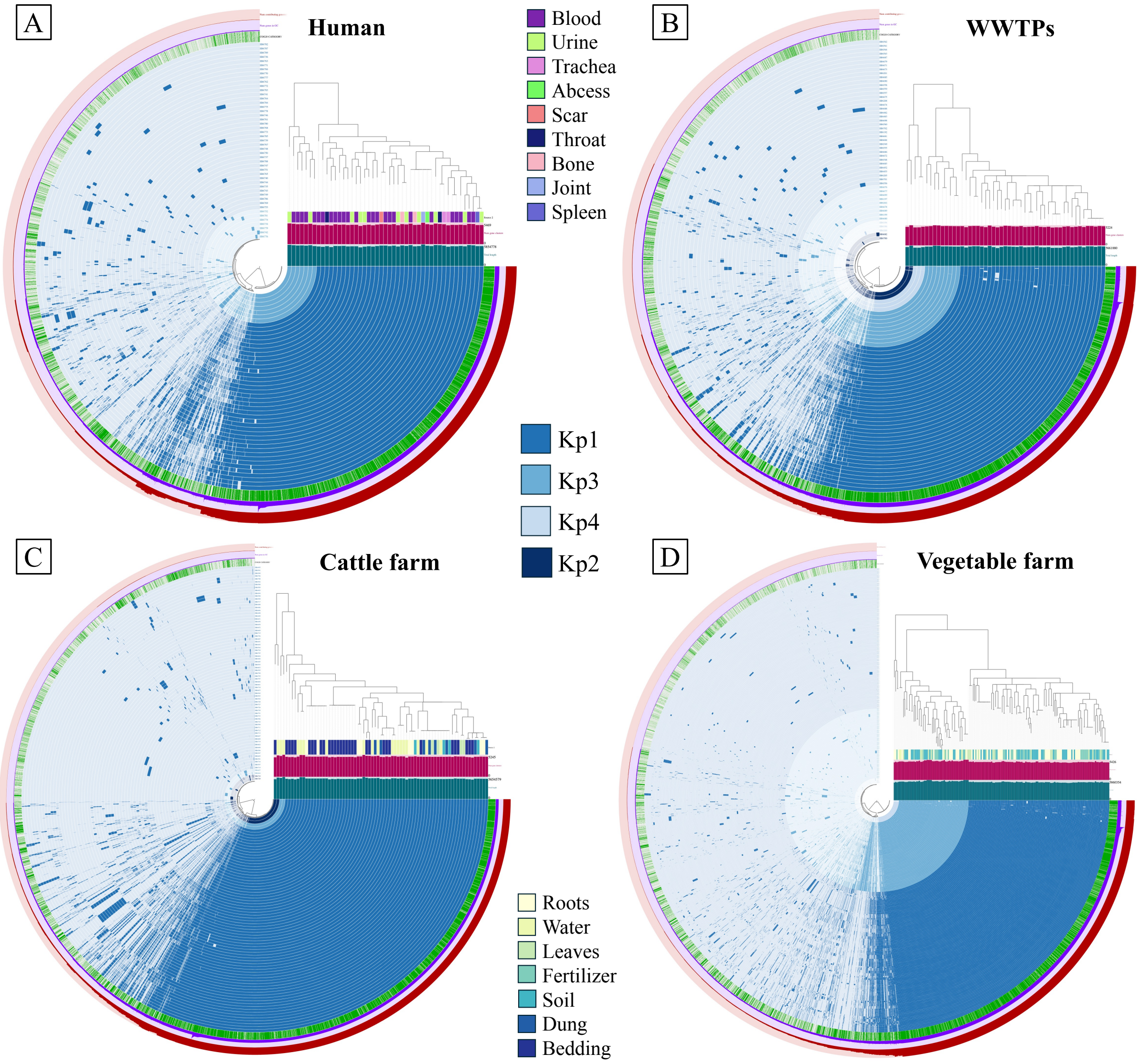


**Figure S5.** Pangenome structure of KpSC populations in each habitat. Each panel displays the gene cluster presence/absence profiles of all KpSC genomes from the same habitat (blue concentric rings). The blue color of the ring represents the KpSC phylogroup (central legend), whereas the blue tone indicates the presence (darker color) and absence (lighter color) of all gene clusters in the genome. The genomes are ordered according to the core-genome SNP tree of the niche population (top-right corner). Between the tree and the rings, from bottom to top, the total genome length (dark green bar plots), the number of genes per genome (maroon bar plots), and the ecological source (top central legend for clinical isolates and bottom central legend for cattle and organic vegetable farms) are indicated for each genome. The tree in the center of the concentric rings represents the presence/absence distribution of each gene cluster across all genomes. The three most outer rings show, from inside to outside, the assignation of gene clusters to a COG category (dark green for assigned COG, light green for unknown COG), the number of genes in each gene cluster (purple ring), and the number of genomes possessing each gene cluster (red ring). **A:** Pangenome structure of human KpSC population. **B:** Pangenome structure of WWTP (wastewater treatment plant) KpSC population. **C:** Pangenome structure of cattle farm KpSC population. **D:** Pangenome structure of organic vegetable farm KpSC population.

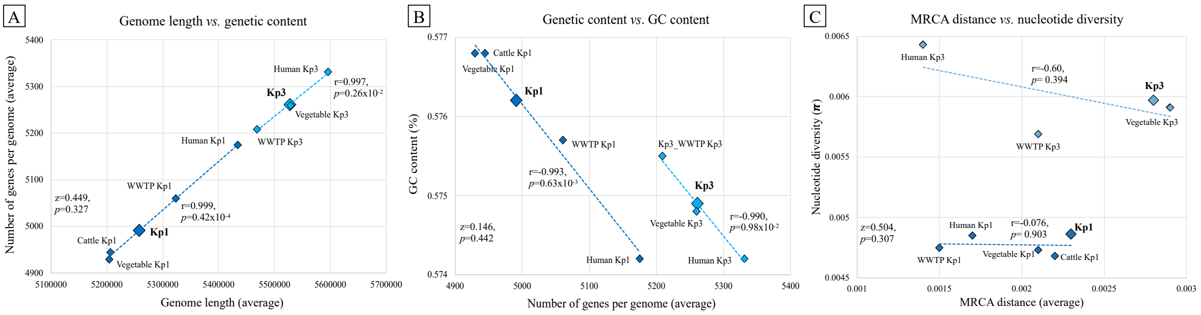


**Figure S6.** Core-genome distances and genome metrics of total and habitat-specific Kp1 and Kp3 populations. Linear fits for Kp1 and Kp3 populations are shown by blue dashed lines, indicating the Pearson's r and *p* values for each population correlation. Fisher's z values for differences of correlations, and their corresponding *p* values, are indicated in the bottom-left corner of each panel. **A:** Association between the genome length and number of genes per genome. **B:** Association between the number of genes per genome and GC content. **C:** Association between the most recent common ancestor (MRCA) distance and nucleotide diversity (π).

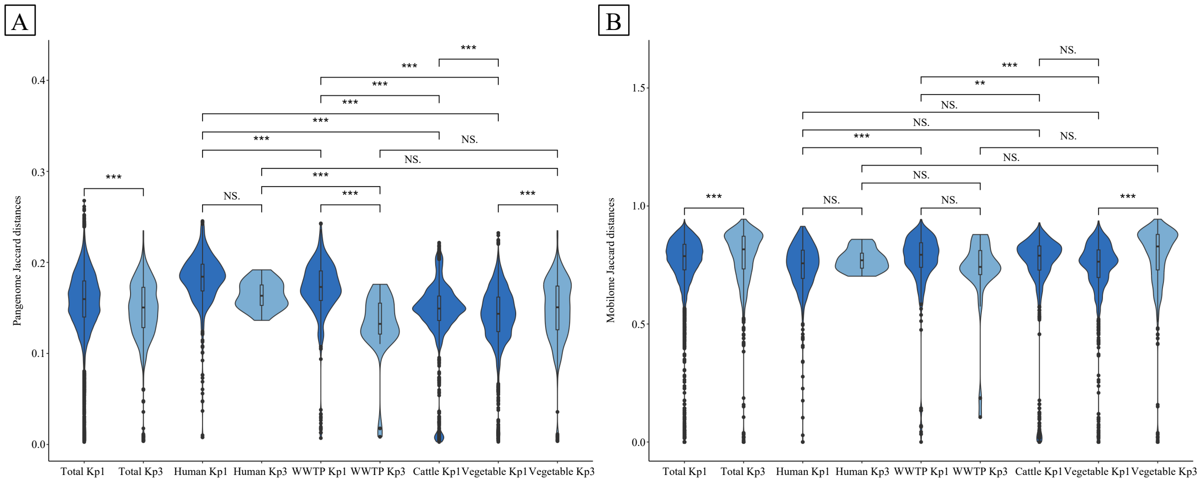


**Figure S7.** Violin plots of pangenome and mobilome diversities of total and niche-specific Kp1 and Kp3 populations. Violin plots show medians and quartiles for Jaccard distances, comparing them by ecological niche and phylogroup. Statistical significance of total Kp1 and Kp3 comparisons was assessed by T-test, whereas the distances between niche-specific populations were evaluated by ANOVA and Tukey's HSD tests. Highly significant differences are indicated by ***. Non-significant differences are indicated by NS. Note the absence of the cattle farm Kp3 population due to the limited number of Kp3 genomes from this niche (n=2). **A:** Pangenome diversity according to Jaccard distances of the total gene cluster content. **B:** Mobilome diversity according to Jaccard distances of the gene clusters belonging to COG annotations linked to mobilome elements (prophages, transposons, plasmids).

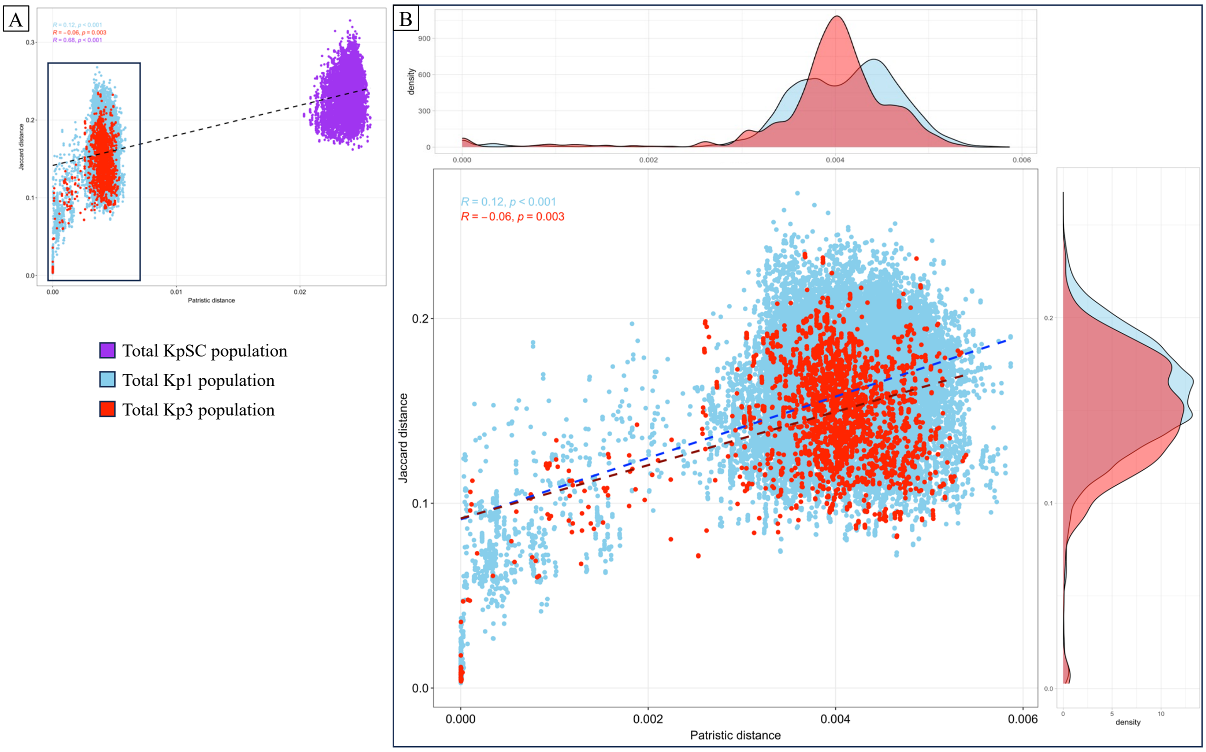


**Figure S8.** Pairwise comparisons between patristic and pangenome Jaccard distances of total KpSC, Kp1 and Kp3 genomes. **A:** Association between patristic and pangenome Jaccard distances of total KpSC genomes. Black dashed line indicates the Spearman r (r=0.68, *p*<0.001). There is a significant positive correlation between patristic distance and pangenome diversity in the total KpSC population. **B:** Zoom-in of the association between patristic and pangenome Jaccard distances of Kp1 and Kp3 genomes. Dark blue dashed line shows the Spearman's r for Kp1 (r=0.11, *p*<0.001). There is a weak positive correlation between patristic distance and pangenome diversity in Kp1 population. Maroon dashed line shows the Spearman's r for Kp3 (r=-0.06, *p*=0.003). There is a weak negative correlation between patristic distance and pangenome diversity in Kp3 population. The Fisher's z-test for Kp1-Kp3 correlation comparison showed a non-significant difference between these two populations (z=1.329, *p*=0.092). Top and side panels show the density of occurrences regarding the patristic distance (top) and pangenome Jaccard distance (right side).

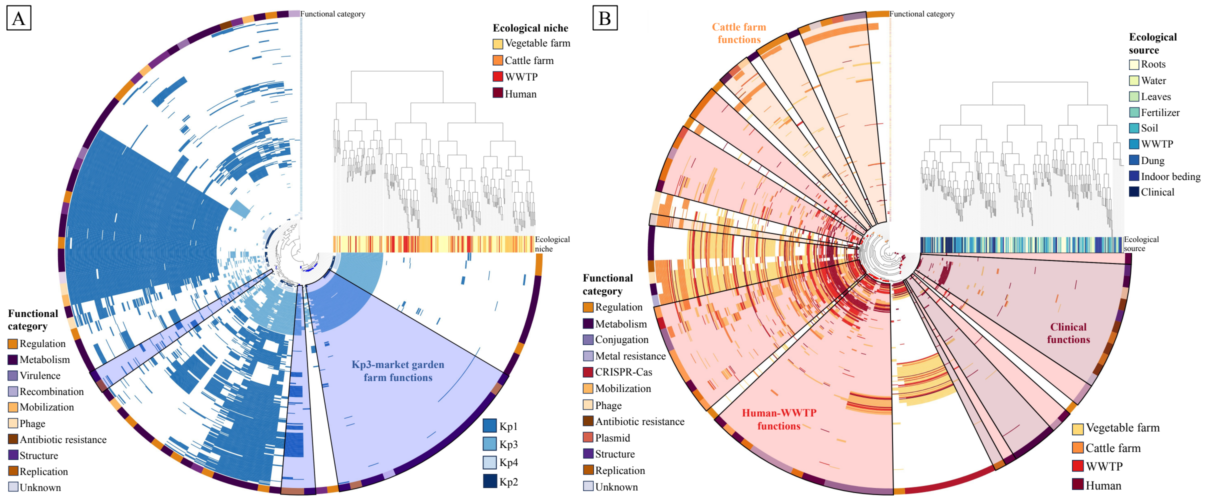


**Figure S9.** Phylogenetically and ecologically enriched functions. Each panel displays the presence/absence profiles of enriched functions in all KpSC genomes (concentric rings). **A:** Phylogenetically enriched functions according to KpSC phylogroups. The rings represent the presence (blue) and absence (white) of phylogenetically enriched functions in each genome, whereas the blue color of the ring indicates the KpSC phylogroup (bottom-right legend). Blue-shaded sectors show the Kp3 phylogenetically enriched functions for the organic vegetable farm environment. The genomes are ordered according to the presence/absence distribution tree of phylogenetically enriched functions (top-right corner). Between the tree and the rings, the ecological niches are indicated for each genome (top-right legend). The tree in the center of the concentric rings represents the presence/absence distribution of each phylogenetically enriched function across all genomes. The most outer ring shows the functional category of all phylogenetically enriched functions (bottom-left legend). **B:** Ecologically enriched functions according to ecological niches. The rings represent the presence (red/orange) and absence (white) of phylogenetically enriched functions in each genome, whereas the red/orange color of the ring indicates the ecological niche (bottom-right legend). Red/orange shaded sectors show the ecologically enriched functions in the different ecological niches according to the labels (bottom-right legend). The genomes are ordered according to the presence/absence distribution tree of ecologically enriched functions (top-right corner). Between the tree and the rings, the ecological sources are indicated for each genome (top-right legend). The tree in the center of the concentric rings represents the presence/absence distribution of each ecologically enriched function across all genomes. The most outer ring shows the functional category of all ecologically enriched functions (bottom-left legend).

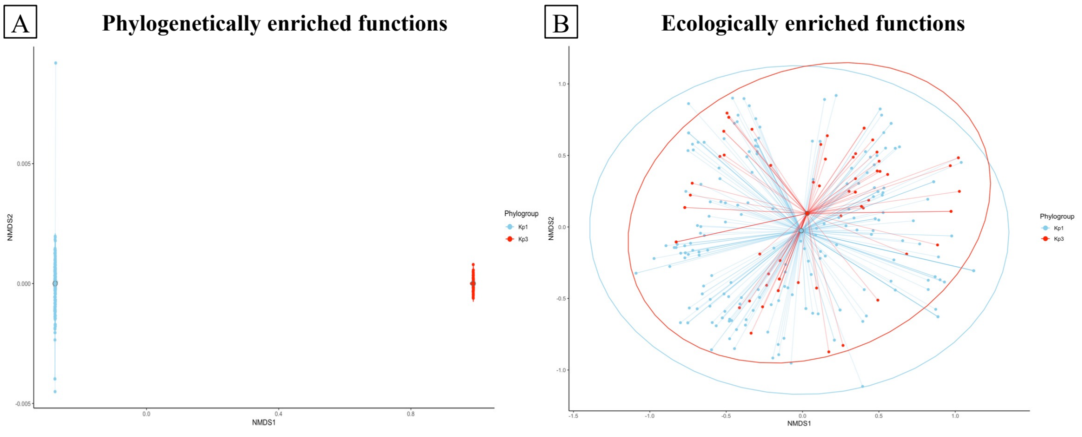


**Figure S10.** Non-metric multidimensional scaling (NMDS) plots of phylogenetically and ecologically enriched functions of total Kp1 and Kp3 populations. NMDS graphs show the divergence of functional content according to Jaccard distances and phylogroups (left side legends), indicating the centroids for the means of each group and clustering them by colored ellipses. **A:** Differential functional content based on presence/absence profiles of phylogenetically enriched functions. **B:** Differential functional content based on presence/absence profiles of ecologically enriched functions.

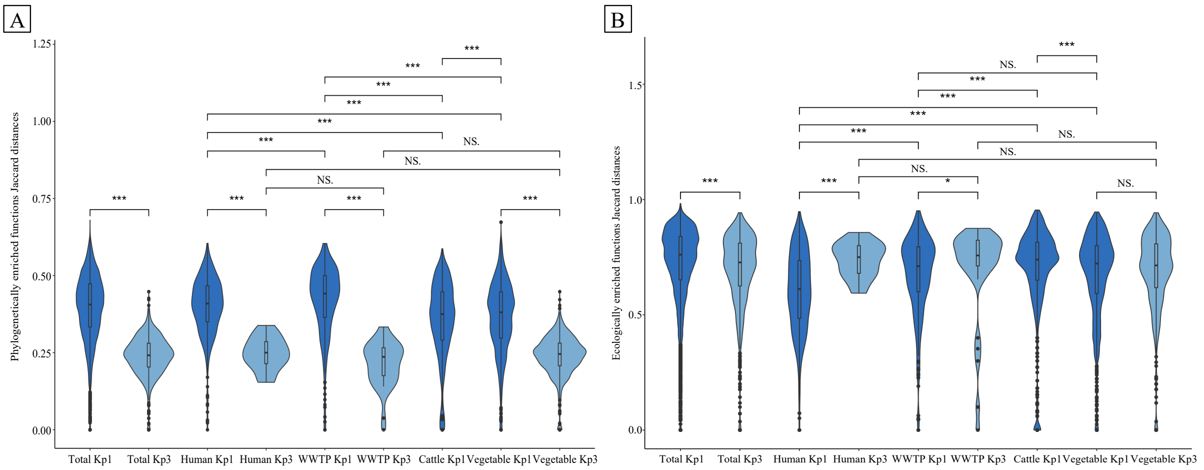


**Figure S11.** Violin plots of functional diversity of phylogenetically and ecologically enriched functions. Violin plots show medians and quartiles for Jaccard distances, comparing them by ecological niche and phylogroup. Statistical significance of total Kp1 and Kp3 comparisons was assessed by T-test, whereas the distances between niche-specific populations were evaluated by ANOVA and Tukey's HSD tests. Highly significant differences are indicated by ***. Non-significant differences are indicated by NS. Note the absence of the cattle farm Kp3 population due to the limited number of Kp3 genomes from this niche (n=2). **A:** Phylogenetically enriched function diversity of total and niche-specific Kp1 and Kp3 populations. **B:** Ecologically enriched function diversity of total and niche-specific Kp1 and Kp3 populations.

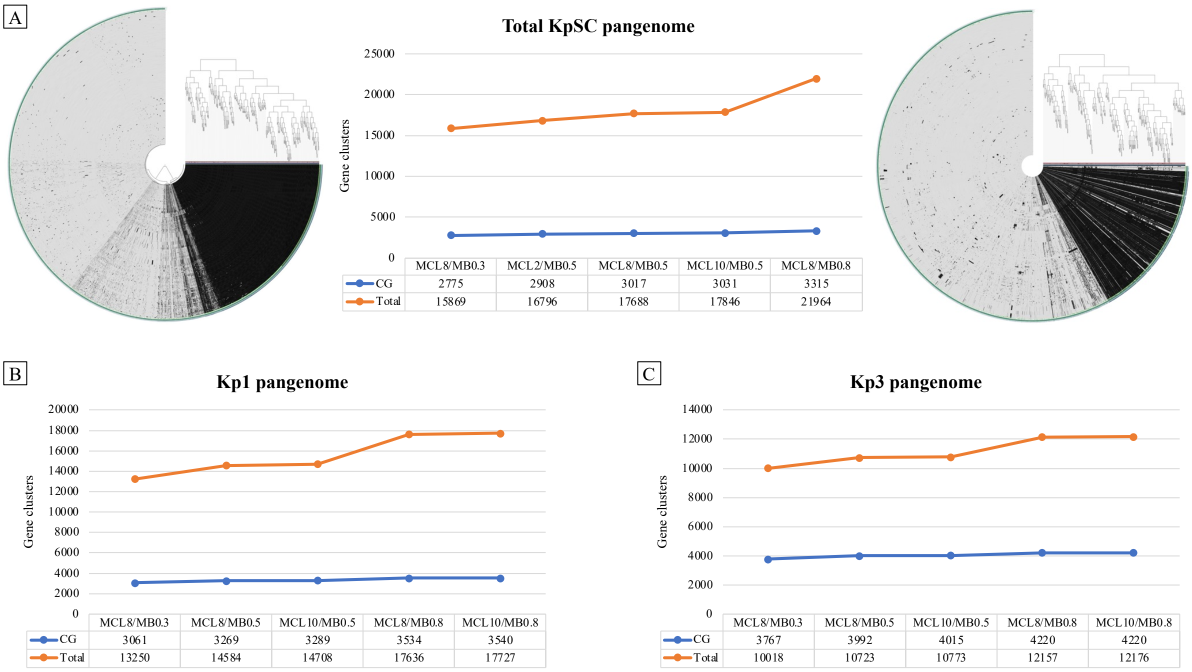


**Figure S12.** Core-pangenome profiles according to different combinations of *minbit* scores (MB) and MCL inflations (MCL). CG: core-genome clusters (blue line). Total: total pangenome clusters (orange line). **A.** Total KpSC population pangenome profiles. Central graph represents core-genome and pangenome trends with different MB and MCL combinations for pangenome clusterization. Left figure shows the anvi'o pangenome graph with MCL8/MB0.3 (less restrictive combination evaluated). Right figure shows anvi'o pangenome graph with MCL8/MB0.8 (most restrictive combination evaluated). Top-right corner trees were constructed according to the presence/absence profiles of gene clusters. Black-shaded areas in the inner concentric rings indicate gene cluster presence, whereas grey-shaded areas indicate gene cluster absence. **B:** Pangenome trends of Kp1 population with different MB and MCL combinations for pangenome clusterization. **C:** Pangenome trends of Kp3 population with different MB and MCL combinations for pangenome clusterization.
